# Estimating Mutation Rates Under Heterogeneous Stress Responses

**DOI:** 10.1101/2023.09.05.555499

**Authors:** Lucy Lansch-Justen, Meriem El Karoui, Helen K. Alexander

## Abstract

Exposure to environmental stressors, including certain antibiotics, induces stress responses in bacteria. Some of these responses increase mutagenesis and thus potentially accelerate resistance evolution. Many studies report increased mutation rates under stress, often using the standard experimental approach of fluctuation assays. However, single-cell studies have revealed that many stress responses are heterogeneously expressed in bacterial populations, which existing estimation methods have not yet addressed. We develop a population dynamic model that considers heterogeneous stress responses (subpopulations of cells with the response *off* or *on*) that impact both mutation rate and cell division rate, inspired by the DNA-damage response in *Escherichia coli* (SOS response). We derive the mutant count distribution arising in fluctuation assays under this model and then implement maximum likelihood estimation of the mutation-rate increase specifically associated with the expression of the stress response. Using simulated mutant count data, we show that our inference method allows for accurate and precise estimation of the mutation-rate increase, provided that the induction of the response also reduces the division rate. Moreover, we find that in many cases, our model of heterogeneous stress responses and the standard model with mutant fitness cost reproduce fluctuation assay data equally well, suggesting that separate experiments would be required to identify the true underlying process. In cases where stress responses are heterogeneous, current methods still correctly infer the effective increase in population mean mutation rate, but we provide a novel method to infer heterogeneity in stress-induced mutation rates.

**Author summary:** How does environmental stress, especially from antibiotics, affect mutation rates in bacteria? This question has often been examined by estimating mutation rates using fluctuation assays, a standard experiment dating back to Luria and Delbrück in the 1940s. We address a gap in currently available estimation methods: we consider cell-to-cell variation in stress responses within bacterial populations, as revealed by recent single-cell studies. Our approach involves a population dynamic model inspired by the DNA-damage response in *E. coli* (SOS response). It accounts for a subpopulation with high expression of the stress response, which increases the mutation rate and decreases the division rate of a cell. Notably, we find that existing models and our heterogeneous-response model reproduce simulated fluctuation assay data equally well in certain scenarios. This emphasises the need for separate experiments to uncover the true underlying processes. Nevertheless, when stress responses are known to be heterogeneous, our study offers a novel method for accurately estimating mutation rates specifically associated with the high expression of the stress response. Uncovering the heterogeneity in stress-induced mutation rates could be important for predicting the evolution of antibiotic resistance.

## 1 Introduction

Bacteria are commonly exposed to adverse conditions, such as starvation, sub-optimal temperatures or toxins, including antibiotics. To cope with these conditions, bacteria have evolved a range of stress responses that enhance viability under stress, often at the expense of a lower growth rate. Some of these response pathways also increase mutagenic mechanisms by, for example, increasing the expression of error-prone polymerases or down-regulating error-correcting enzymes [1, 2]. It has been proposed that this so-called ‘stress-induced mutagenesis’ in bacterial cells could accelerate the evolution of populations that are poorly adapted to their environment [3, 4]. Consequently, inhibiting bacterial stress responses has been suggested to prevent antibiotic resistance evolution, and gained some experimental support [5, 6].

Several studies report increased mutation rates in bacterial populations exposed to sublethal antibiotic concentrations [7, 8, 9, 10, 11, 12, 13]. Typically, these mutation rates have been measured with fluctuation assays [14, 15]. The standard experiment involves growing parallel cultures under permissive conditions while mutations occur randomly and subsequently plating the cultures onto selective plates. The mutation rate is estimated from the distribution of the number of mutants on the plates. To quantify the mutationrate increase associated with stress, the standard experiment is repeated, but with the cultures exposed to a stressor during the growth phase. Then, the *stress-induced* mutation rate is estimated and compared with the mutation rate under permissive conditions. However, stress impacts the growth of bacterial cells in several ways, which are neglected in commonly applied estimation methods, potentially leading to biased estimates of the mutation rate. For instance, increased cell death leads to overestimating the mutation rate [16]. Another effect that has not yet been addressed is cell-to-cell heterogeneity in stress responses.

In recent years, single-cell experiments have revealed extensive heterogeneity in the expression of stress responses in bacterial populations [17, 18, 19, 20, 21, 22, 23, 24, 25, 26]. Heterogeneity can arise for various reasons, including stochastic expression of genes involved in stress responses, especially where the corresponding proteins are initially present in small numbers [18, 19, 20], phenotypic variability in the stability of key regulators [23], or micro-environmental variation in cell-to-cell interactions [26]. Positive and negative feedback loops are common features of stress response regulatory networks, which can generate, amongst other features, cell-to-cell variation [27]. In some cases, a subpopulation of cells showing elevated stress responses has been directly associated with a higher rate of DNA mismatches or higher mutant frequency [18, 19, 20, 22, 13, 24].

In addition to mutagenic mechanisms, stress responses can alter cell division and death rates. For example, the widely studied SOS response, which leads to the transcriptional induction of approximately 40 genes after exposure to DNA damage, involves inhibition of cell division and induction of error-prone polymerases that could increase mutation rate [28, 29]. Single-cell studies using fluorescent reporters for the SOS response in *E. coli* have revealed that its expression is highly heterogeneous and that this heterogeneity can be approximated as a bimodal response, with some cells expressing low levels of SOS under DNA damage whilst others show a very high level of expression [17, 19, 25]. Overall, heterogeneously expressed stress responses are therefore likely to impact both bacterial population dynamics and mutational input during the growth phase of a fluctuation assay, and it is unclear whether estimation methods that neglect heterogeneity in stress responses produce reliable results.

In this study, we present a population dynamics model that considers cell-to-cell heterogeneity in stress responses. Motivated by the SOS response, we describe two discrete subpopulations of cells, where high expression of the stress response is associated with both a higher mutation rate and a lower division rate than in cells with low expression. We derive the resulting mutant count distribution in the total population and implement maximum likelihood estimation of the mutation-rate increase associated with the induction of the stress response. We test the performance of our method using stochastic simulations of fluctuation assays under permissive and stressful conditions, including robustness to biologically realistic model deviations such as mutant fitness costs and cell death. We also apply formal model comparison to assess whether cell-to-cell heterogeneity could be detected from fluctuation assays alone.

## 2 Model & Methods

### 2.1 Population dynamics with heterogeneous stress responses

We model the population dynamics of bacterial cultures during the growth phase of a fluctuation assay, (i) under permissive conditions and (ii) under exposure to some form of stress that induces a mutagenic stress response in the cells. For the permissive conditions, we assume the standard model: an exponentially growing population without cell death or differential mutant fitness. For growth under stressful conditions, our model considers cellto-cell heterogeneity in the expression of the stress response. In particular, for simplicity, we assume that under stress, the population can be divided into two subpopulations: one with a low expression level of the stress response (here referred to as response switched *off*, even if strictly speaking the response is not fully off but very low) and the other with a high expression level (here referred to as response switched *on*). This *heterogeneous-response model* differs from models used in currently available mutation-rate estimation methods, which assume all cells respond to the stress in the same way (*homogeneous-response model* ).

In the following, we describe the full heterogeneous-response model we use in the simulations to test our inference algorithm, approximations we make to derive the mutant count distributions used in the inference, and the homogeneous-response model used in current inference methods, summarised in Figure 1.

**Figure 1:**
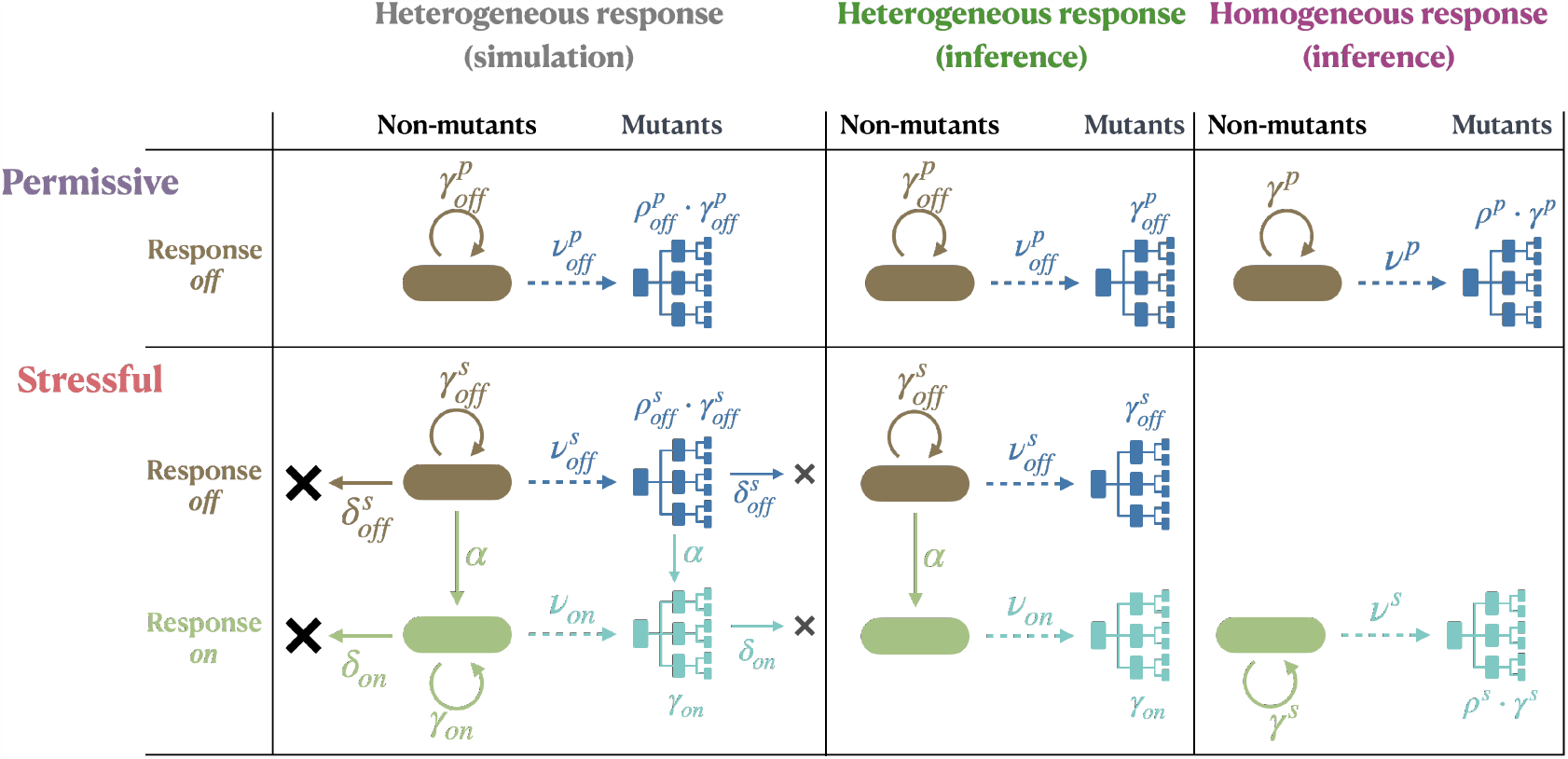
Schematic illustrating the heterogeneous- and homogeneous-response models used in simulations and inference. Cell death (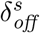 and *δ*_*on*_) and differential mutant fitness (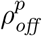 and 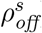) are included only in extended versions of the simulation model where explicitly stated.

#### 2.1.1 Heterogeneous-response model

We consider a bacterial population previously growing under permissive conditions and now exposed to stress. Cells induce a stress response with a constant switching rate *α*, leading to the emergence of a response-*on* subpopulation. Inducing the stress response alters the mutation rate of the cells but potentially also their division and death rates. We assume that, as long as the stress persists, cells do not switch the response *off* again.

The above-described population dynamics result in the following set of coupled differential equations for the population sizes of the non-mutant response-*off* and response-*on* subpopulations, *n*_*off*_ and *n*_*on*_, respectively:

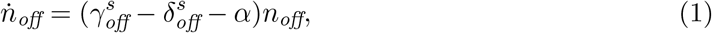

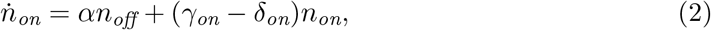

Here, 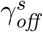 is the division rate of the response-*off* subpopulation under stress (which can be different than under permissive conditions, 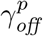) and 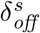 its death rate, and *γ*_*on*_ and *δ*_*on*_ the division and death rate of the response-*on* subpopulation. The solution to these equations is given by

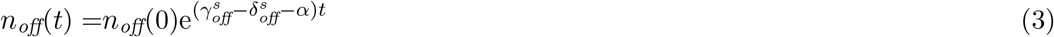

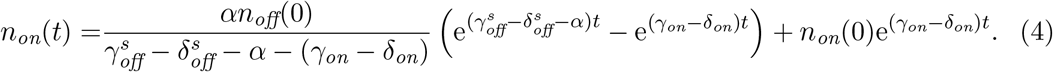

with *n*_*off*_(0) and *n*_*on*_(0) denoting the number of response-*off* and response-*on* cells, respectively, before the exposure to the stress.

This approach assumes that the non-mutants, including the initially small response-*on* subpopulation, can be treated deterministically. We test the validity of this assumption using stochastic simulations: we simulate switching *on* of the response as a time-inhomogeneous Poisson process and the growth dynamics of the response-*on* subpopulation as a stochastic birth-death branching process. Then, we compare the resulting population size with equation 4. We find that deviations from the deterministic prediction are negligible for a wide range of switching rates and division rates of response-*on* cells and for zero and small initial sizes of the response-*on* subpopulation (Supplementary Material Figure S1).

In the simulations to test our inference algorithm, we use equations 3 and 4 for the dynamics of the non-mutant population, setting *n*_*on*_(0) = 0. For the inference itself, however, we approximate equation 4 as

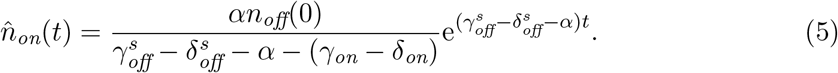

because this allows us to derive an analytical expression for the mutant count distribution later. This approximation is valid when the initial population size of the response-*on* subpopulation is comparably small, *n*_*on*_(0) *≪ n*_*off*_(0), and its growth is slower than the growth of the response-*off* subpopulation,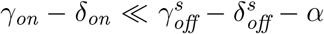. As a consequence of this approximation, the total population grows exponentially with a *population growth rate* of

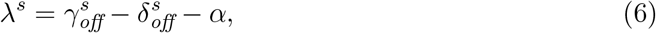

and the response-*on* subpopulation makes up a constant fraction of

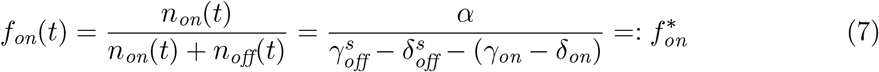

For the inference, we assume that the fraction of the response-*on* subpopulation at the end of the growth phase, *f*_*on*_, is a good approximation of 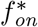.

We define the *relative switching rate* as

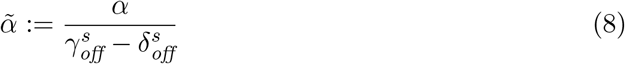

and the *relative fitness* of response-*on* compared to response-*off* cells as

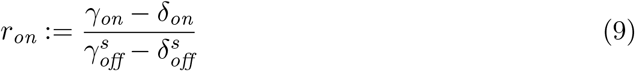

and thereby obtain

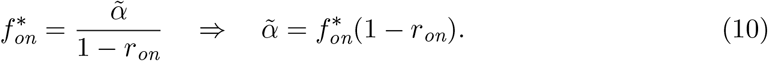

In the simulations and the inference, we consider mutations in the response-*off* and the response-*on* subpopulation to occur according to time-inhomogeneous Poisson processes and treat the dynamics of the resulting mutants stochastically. Mutations arise in each subpopulation at rates 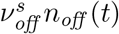 and *ν*_*on*_*n*_*on*_(*t*) (simulation) or 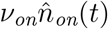 (inference), re-spectively. Note that *ν* describes the mutation rate to the phenotype of interest selected on the plates in the fluctuation assay (mutations per cell per unit time, also called instantaneous mutation rate). In the simulations, we set the duration *t*_*f*_ of the growth phase such that the expected number of *mutations* (not *mutants*) is constant across the considered parameter ranges (Supplementary Material section **S2**).

In a fluctuation assay, the selective agent on the plates is normally chosen to be unrelated to the stressor applied in the growth phase (e.g. two different antibiotics with no cross-resistance). Therefore, we assume that mutation itself does not alter the stress response, and mutants can induce the response equivalently to non-mutants. Nonetheless, mutations might affect the fitness during the growth phase, which we implement by an altered division rate. Together, these assumptions result in a two-type branching process describing the mutant response-*off* and response-*on* subpopulations, with respective birth rates 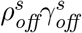 and *ρ*_*on*_*γ*_*on*_, respective death rates 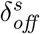 and *δ*_*on*_, and switching at rate *α*.

When testing our inference algorithm, we simulate the mutant dynamics using the two-type branching process described above. For the inference, however, we neglect the induction of the stress response in the mutants and assume that mutations have no fitness effect. For mathematical convenience, we consider switching as a reduction in the division rate of response-*off* mutants by *α* instead (birth rate equal to 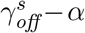). This approximation allows us to derive an analytical expression for the mutant count distribution because the dynamics of response-*off* and response-*on* mutants are now independent birth-death branching processes and the total mutant count distribution is given by the sum of the mutant contribution of each subpopulation as two independent random variables.

The complete documentation of all population dynamic functions can be found in the file called **population dynamics.jl** at https://github.com/LucyL-J/Quantifying-SIM.

#### 2.1.2 Mutant count distributions

We use the results from [30], who derive the mutant count distribution for the general case of an exponentially growing population with initial population size *N*_*i*_, population growth rate *λ*, mutation rate *ν*, mutants with birth rate *b* and death rate *d*, and 100% plating efficiency. They calculate the probability-generating function *G*(*z*) of the mutant count distribution when the population reaches a final population size *N*_*f*_ . For *N*_*f*_ *≫ N*_*i*_ (neglecting initial population size effects), and defining the per-generation mutation rate as 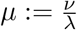, the differential fitness of mutants as 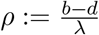, and the extinction probability of mutants as 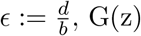, G(z) is given by

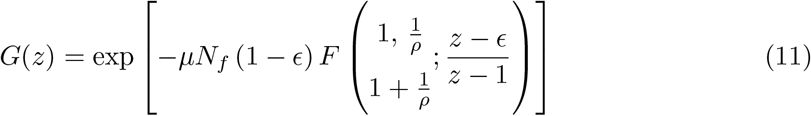

with *F* being the hypergeometric function. Moreover, the authors [30] give a recursive expression for the probabilities *p*_*m*_ to observe *m* mutants, which allows for the numerically relatively inexpensive calculation of the likelihood of observing empirical mutant count data.

Using equation 11, we can substitute the corresponding parameters to obtain the mutant count distributions under both permissive and stressful conditions for the heterogeneousresponse model. Importantly, this approach assumes that all mutants, including response*on* cells, form colonies on the selective plates.

We assume that under permissive conditions, there is no response-*on* subpopulation and consider the simple case that mutants have the same division rate as non-mutants (*ρ* = 1). This results in the following mutant count distribution under permissive conditions:

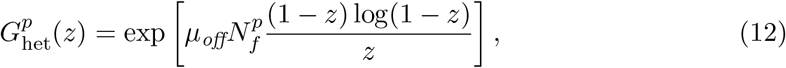

where 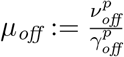 describes the *per-division* mutation rate, which equals the *per-generation* rate as, under permissive conditions, the population growth is solely determined by cell division.

Under stressful conditions, on the other hand, the population growth rate depends on the switching rate *α >* 0 via

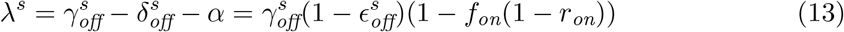

which we have re-written as a function of the division rate 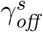 and the extinction probability 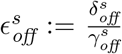 of response-*off* cells, and the fraction *f*_*on*_ and relative fitness *r*_*on*_ of the response-*on* subpopulation. Importantly, we assume that the *per-division* mutation rate of response-*off* cells is the same under stressful as under permissive conditions, 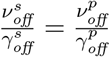.Then, the per-generation mutation rates of response-*off* and response-*on* cells are given by

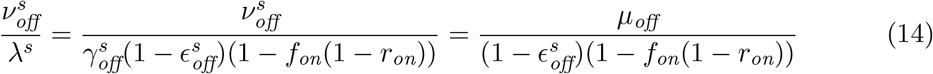

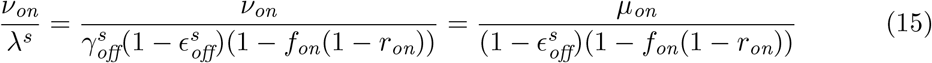

with 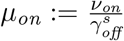 . For response-*on* cells, the relative fitness of mutants *ρ*_*on*_ (relative to the population growth rate) can be expressed via

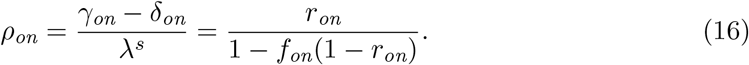

The total mutant count is the sum of the independent contributions from the response-*off* and response-*on* subpopulations. Therefore, the probability generating function of the total mutant count under stressful conditions, 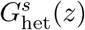, is given by the product 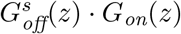, where

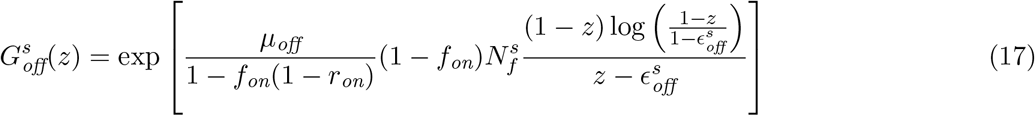

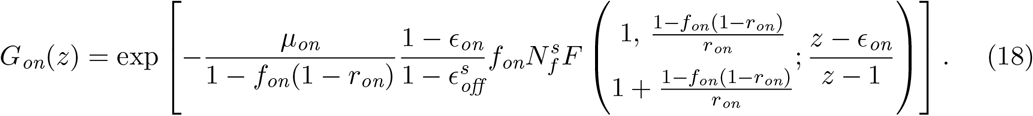

In the case of *γ*_*on*_ = 0, the contribution to the mutant count from the response-*on* subpopulation follows a Poisson distribution and equation 18 reduces to

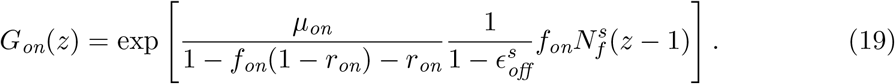

Note that in this case 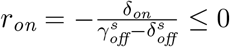.

The homogeneous-response model assumes that all cells respond to the stress similarly. The stress response results in a change in the per-generation mutation rate, *µ*^*p*^ *→ µ*^*s*^, and potentially the differential fitness of mutants compared to non-mutants, *ρ*^*p*^ *→ ρ*^*s*^. Therefore, the probability-generating functions under permissive and stressful conditions are given by

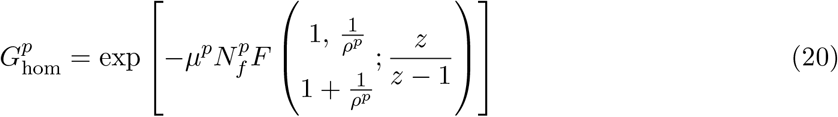

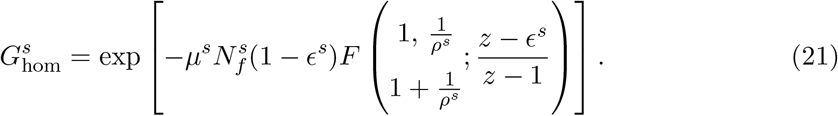

### 2.2 Estimation of the mutation-rate increase under stress

We use joint maximum likelihood estimation to find the set of model parameters for which the empirically observed mutant count data are most likely. We consider the permissive and the stressful conditions concurrently and jointly infer the respective parameters. The joint inference crucially relies on our assumption that, even though the division rate might change under stress, the *per-division* mutation rate of response-*off* cells is the same under permissive and stressful conditions.

We define a log-likelihood function

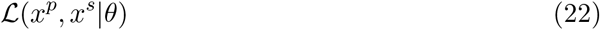

that is a measure for the probability of observing the mutant counts *x*^*p*^ and *x*^*s*^ under permissive and stressful conditions, respectively, for a given model and model parameters *θ*. It is calculated from the mutant count distributions described above, using the results of [30] to compute the probabilities recursively from the probability generating functions. Then, we use an optimisation algorithm implemented in the Julia [31] package Optim.jl (https://julianlsolvers.github.io/Optim.jl/stable/), to find the parameters that maximise this log-likelihood function. The initial parameters for the optimisation are calculated using, among others, the generating function method described in [32].

#### 2.2.1 Inferred and measured parameters

The mutant count distributions under permissive and stressful conditions derived for the heterogeneous-response model, equations 12, 17 and 18, depend on the following parameters, some of which are inferred by our algorithm and some of which must be set to fixed values.

- *µ*_*off*_: The per-division mutation rate of response-*off* cells is inferred.
- *µ*_*on*_: The per-division mutation rate of response-*on* cells is also inferred. Note that this rate is given in ‘mutations in response-*on* cells per divisions in response-*off* cells’ because the division rate of response-*on* can be zero. This definition makes the mutation rates *µ*_*off*_ and *µ*_*on*_ directly comparable.
- 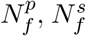: The final population sizes under permissive and stressful conditions are taken as known quantities in our inference and would normally be measured separately, typically through plating a few cultures on non-selective media and colony counting.
- *r*_*on*_: The relative fitness of response-*on* compared to response-*off* cells is an optional inference parameter. By default, it is set to *r*_*on*_ = 0, but it can also be inferred (Results section 3.3) or set to another value if an estimate of *r*_*on*_ is available (Supplementary Material Figure S4).
- *f*_*on*_: The fraction of the response-*on* subpopulation at the end of the growth phase is either taken as a known quantity, which would have to be measured in a separate experiment (e.g. microscopy or flow cytometry), or inferred (Results section 3.4).
- 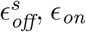: The extinction probabilities of response-*off* cells under stress and of response*on* cells are set to zero in the inference, corresponding to no cell death. Neglecting cell death is in common with most existing approaches, but see [16]. We test the robustness of this assumption in Results section 3.2.

Ultimately, we are interested in quantifying the relative increase in mutation rate associated with induction of the stress response, that is, 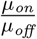 . In addition, we calculate the (relative) increase in *population mean mutation rate*

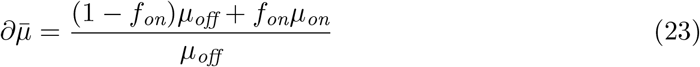

which is directly comparable to inference results from the homogeneous-response model. Note that, strictly speaking, the fraction of the response-*on* subpopulation changes with time until the stationary fraction 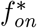 is reached. For simplicity, we use the fraction *f*_*on*_ at the end of the growth phase to define the increase in population mean mutation rate 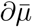 (equation 23).

The mutant count distributions under permissive and stressful conditions derived for the homogeneous-response model, equations 20 and 21, depend on the following parameters:

- *µ*^*p*^, *µ*^*s*^: The per-generation mutation rates under permissive and stressful conditions are inferred.
- 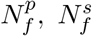: The final population sizes are, as for the heterogeneous-response model, taken as known quantities.
- *ρ*^*p*^, *ρ*^*s*^: The differential fitness of mutants compared to non-mutants is an optional inference parameter, also previously implemented, for example, in [33]. By default, it is set to *ρ*^*p,s*^ = 1. Alternatively, we take *ρ*^*p*^ = *ρ*^*s*^ as a joint inference parameter, or *ρ*^*p*^ and *ρ*^*s*^ as two separate inference parameters.
- *ϵ*: The extinction probability is set to zero in the inference because we neglect cell death, as above.

Ultimately, the quantity of interest is the mutation-rate increase associated with the stress, that is, 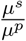. Note, however, that both *µ*^*p*^ and *µ*^*s*^ are population-wide rates and 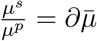 is equivalent to the (relative) increase in population mean mutation rate under stress.

The complete documentation of all inference algorithms can be found in the file called **inference.jl** at https://github.com/LucyL-J/Quantifying-SIM.

### 2.3 Testing the inference algorithm using stochastic simulations

To test the inference algorithm, we perform stochastic simulations of pairs of fluctuation assays, one under permissive and the other under stressful conditions. Then, we infer the respective parameters under heterogeneous- and homogeneous-response models and compare them to the true underlying parameters; see Figure 1 for schematics of the models used in simulation and inference. For each parameter set used, we simulate *R* = 100 pairs of fluctuation assays, with *c* = 50 parallel cultures per fluctuation assay. To plot the estimated parameters across the *R* = 100 simulations, we use boxplots, where each box shows the median and interquartile range with whiskers extending to 1.5 times the interquartile range and any outliers outside that range represented as dots.

#### 2.3.1 Accuracy and precision

First, we evaluate the estimation of the mutation-rate increase associated with the stress response. For this purpose, we simulate fluctuation assays under the heterogeneous-response model without cell death or differential mutant fitness, and for (a) *γ*_*on*_ = 0, with varying *{µ*_*on*_, *α}* (Results section 3.1), and (b) varying *γ*_*on*_ *>* 0 (Results section 3.3). Then, we infer the underlying model parameters under the same model as simulated (with the only difference that switching of mutants and initial population size effects are neglected in the inference, as described above). In case (a), we set the relative fitness of response-*on* cells, *r*_*on*_, to zero, and in case (b), we consider both setting *r*_*on*_ = 0 and inferring *r*_*on*_. In either case, we assume that separate estimates of 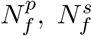, and *f*_*on*_ are available, and we take these as fixed values.

We evaluate the accuracy of the estimation by calculating the relative error of the median across the *R* replicates,

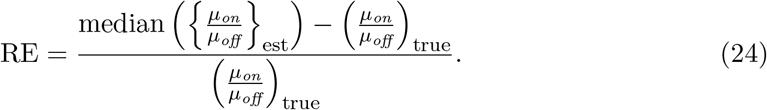

Here, a positive or negative relative error implies over- or underestimation, respectively. We also evaluate the precision of the estimation by calculating the coefficient of variation across the *R* replicates,

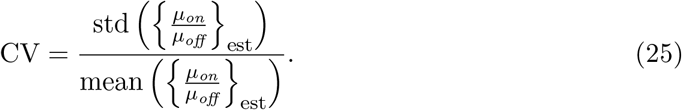

mean where ‘std’ denotes standard deviation.

#### 2.3.2 Robustness to model deviations

Next, we evaluate the robustness of the estimation to model deviations. For this purpose, we simulate fluctuation assays under a more complex version of the heterogeneous-response model than is used in the inference. We simulate with (a) cell death in three different forms: 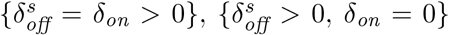, or 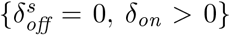 (Results section 3.2 and Supplementary Material section **S3**); and (b) mutants having a different fitness than non-mutants (Supplementary Material section **S4**). In the inference, however, we neglect these effects and evaluate whether this introduces any biases to the estimates. Again, we assume that separate estimates of 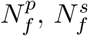, and *f*_*on*_ are available.

#### 2.3.3 Model selection: heterogeneous versus homogeneous response

Finally, we evaluate whether it is possible to identify the heterogeneity of stress responses from mutant count data alone. In this case, we suppose we do not have separate experimental data showing heterogeneity and, therefore, do not have an estimate of *f*_*on*_. For this purpose, we simulate fluctuation assays under the heterogeneous-response model for varying *γ*_*on*_. Then, we infer the respective model parameters assuming (a) the heterogeneousresponse model with either *r*_*on*_ = 0 or *r*_*on*_ inferred, as well as *f*_*on*_ as an inference parameter; and (b) the homogeneous-response model with either *ρ*^*p*^ = *ρ*^*s*^ = 1 as a fixed value, *ρ*^*p*^ and *ρ*^*s*^ inferred individually (Results section 3.4), or *ρ*^*p*^ = *ρ*^*s*^ inferred jointly (Supplementary Material section **S6**). To decide which model fits the observed mutant count data best, we use the Akaike information criterion (AIC), defined as

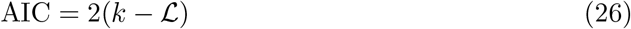

where *k* is the number of inferred parameters of the model. We select the model with the lowest AIC. However, if the difference in AIC between the (a) best-fitting heterogeneousand (b) the best-fitting homogeneous-response model is within *±*2, we say that neither of the models can be selected because their AICs are comparable.

We also evaluate the models’ performances in estimating the relative increase in population mean mutation rate. In case (a), the population mean mutation rate increase is calculated as 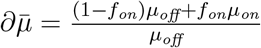 and in (b) as 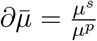 .

The complete documentation of the computational analysis can be found in the **README** at https://github.com/LucyL-J/Quantifying-SIM.

## 3 Results

We aim to estimate the increase in mutation rate associated with the induction of the stress response when this response is heterogeneously expressed across the bacterial population. In particular, we consider cases in which the population can be divided into two discrete subpopulations: one with a low expression level (response *off* ) and the other with a high expression level (response *on*). The key principle of our method is to jointly infer their mutation rates from mutant count data obtained from a pair of fluctuation assays under permissive and stressful conditions. For the latter, we need to disentangle the contributions from the response-*off* and response-*on* subpopulations. The success of this method relies on the changing shape of the mutant count distribution under stress, which occurs if there is a highly mutating but slowly dividing response-*on* subpopulation.

To evaluate the performance of our method, we use simulated mutant count data to compare the estimated parameters with the true underlying ones. First, we explore how the accuracy and precision of our method depend on the model parameters by simulating and inferring under the same model. Then, we test the robustness of our method to model deviations by simulating under a more complex model than used in the inference. Finally, we determine under what conditions the heterogeneous-response model can be distinguished from the homogeneous-response model assumed in currently available methods by inferring under both models and comparing how well they fit simulated data.

In all simulations, we set the initial population size to 10^4^ and the initial fraction of the response-*on* subpopulation to zero. Moreover, we consider the duration of the growth phase such that the expected number of mutations equals one. This way, the resulting number of resistant mutant colonies on each selective plate is usually within an experimentally countable range of zero to a couple hundred (Supplementary Material section **S2**). Table 1 summarises the default parameters used in the simulations, while parameters that vary are specified in the relevant Results section. For each parameter combination, we simulate *R* = 100 pairs of fluctuation assays under permissive and stressful conditions, with *c* = 50 parallel cultures per assay. Generally, we assume that the final population sizes 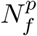 and 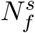 and the fraction of the response-*on* subpopulation *f*_*on*_ are known from separate experimental measurements, except for Results section 3.4 where we infer without an estimate of *f*_*on*_.

**Table 1:** Parameters used in the simulations unless explicitly specified otherwise. For simplicity, we set the division rate under permissive and stressful conditions to 1 *h*^*−*1^. The switching rate for the SOS response in *E. coli* has been estimated using single-cell imaging [25]. The mutation rate is based on rifampicin resistance, a selective marker commonly used in fluctuation assays. In *E. coli*, the number of point mutations conferring rifampicin resistance has been estimated to be 79 [34] and the mutation rate between 0.2 ·10^*−*10^ and 5 ·10^*−*10^ nucleotides per generation in permissive conditions [35], yielding a mutation rate of 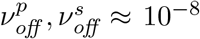 per unit time for response-*off* cells. Meanwhile, the mutation rate under induction of the stress response (*ν*_*on*_) is set to a default of 100 times higher, comparable to the increase associated with genetic mutators [36, 35].

| Symbol | Condition | Definition | Numerical value |
| --- | --- | --- | --- |
| $\gamma_{off}^p$ | Permissive | Division rate response-*off* cells | $1 h^{-1}$ |
| $\rho_{off}^p$ | Permissive | Differential mutant fitness response-*off* cells | 1 |
| $\nu_{off}^p$ | Permissive | Mutations per unit time response-*off* cells | $10^{-8} h^{-1}$ |
| $\gamma_{off}^s$ | Stressful | Division rate response-*off* cells | $1 h^{-1}$ |
| $\rho_{off}^s$ | Stressful | Differential mutant fitness response-*off* cells | 1 |
| $\nu_{off}^s$ | Stressful | Mutations per unit time response-*off* cells | $10^{-8} h^{-1}$ |
| $\delta_{off}^s$ | Stressful | Death rate response-*off* cells | $0 h^{-1}$ |
| $\gamma_{on}$ | Stressful | Division rate response-*on* cells | $0 h^{-1}$ |
| $\rho_{on}$ | Stressful | Differential mutant fitness response-*on* cells | 1 |
| $\nu_{on}$ | Stressful | Mutations per unit time response-*on* cells | $10^{-6} h^{-1}$ |
| $\delta_{on}$ | Stressful | Death rate response-*on* cells | $0 h^{-1}$ |
| $\alpha$ | Stressful | Switching rate | $0.05 h^{-1}$ |

### 3.1 Estimating the mutation-rate increase

First, we evaluate our method’s performance in the best-case scenario; that is, we simulate and infer under the same model, beginning with the simplest model of heterogeneous stress responses without cell death, with mutant fitness equal to non-mutant fitness, and with zero division rate of response-*on* cells. We simulate for a range of mutation rates in response-*on* cells, *ν*_*on*_ *∈* [10^*−*5^, 10^*−*8^] *h*^*−*1^ and switching rates *α ∈* [0.001, 0.1] *h*^*−*1^. Note that the per-division rate *µ*_*on*_ and the per-unit-time rate *ν*_*on*_ are equivalent here because we set the division rate 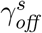to one.

For the inference, we consider the same model as used in the simulations with the only exception that, in the inference, we neglect switching *on* the stress response in mutants and initial population size effects, see Model & Methods. For each set of mutant count data, we infer the mutation rates *µ*_*off*_ and *µ*_*on*_ and calculate the mutation-rate increase 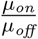. To quantify the accuracy of our method, we calculate the relative error of the median of our estimated mutation-rate increases (equation 24). Additionally, we use the coefficient of variation (equation 25) across the estimates to quantify our method’s precision.

Comparing the estimated with the true mutation-rate increase 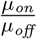, we find that the accuracy and precision improve with increasing 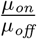 and relative switching rate 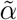 (Figure 2B, 2C). For example, for 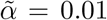, when 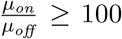, 75% of estimates lie within 2*−*fold of the true mutation-rate increase and the estimation is unbiased (Figure 2A). For smaller 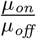, on the other hand, the variation in the estimates becomes large. Nonetheless, if the mutation-rate increase is estimated to be *>* 100, the true increase is very likely to be *>* 25.

**Figure 2:**
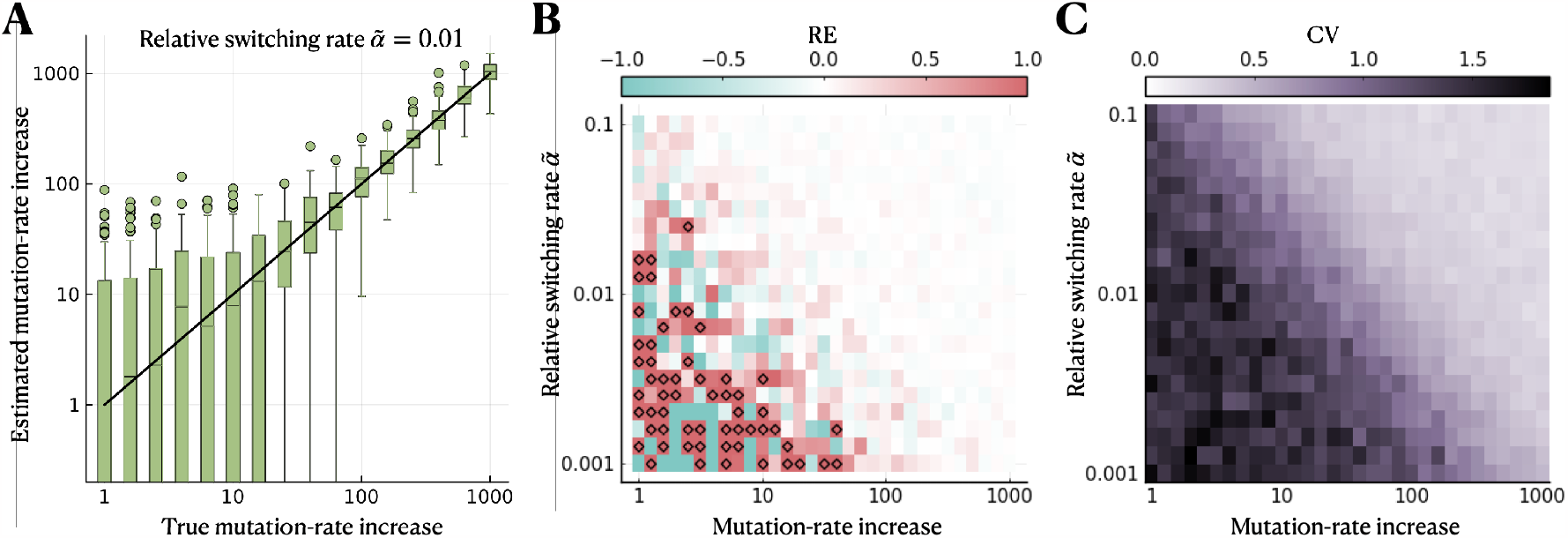
Estimation of the mutation-rate increase is accurate and precise when the heterogeneity in mutation rate is sufficiently large. We simulate using the simplest model of heterogeneous stress responses (without cell death or differential mutant fitness and with zero division rate of response-*on* cells) and infer the mutation rates *µ*_*on*_ and *µ*_*off*_, assuming the same model in the inference. **A** Estimated compared to true mutation-rate increase 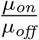 for a range of values of 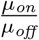 and a relative switching rate of 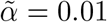. **B** Median relative error of estimated compared to true mutation-rate increase for a range of 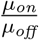 and 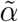. Over/underestimation is shown in red/blue, and diamonds indicate a relative error greater than one. **C** Coefficient of variation across estimates. The parameter ranges used in the simulations are *ν*_*on*_ *∈* [10^*−*5^, 10^*−*8^] *h*^*−*1^ and *α ∈* [0.001, 0.1] *h*^*−*1^.

The composite parameter 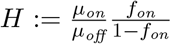, which is a measure for the mutation-rate heterogeneity in the population, determines our method’s performance. This can be seen from the mutant count distribution derived in Model & Methods. For the simplest model considered here, the probability-generating function of the mutant count distribution under stress, obtained from equations 17 and 19 is given by

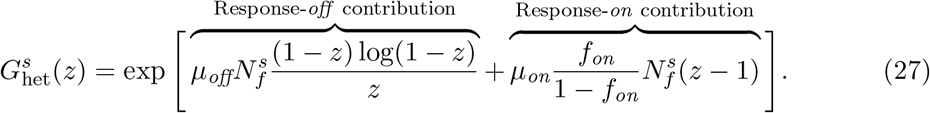

The mutation-rate heterogeneity determines the shape of the distribution as for *H ≫* 1, the contribution of the response-*on* subpopulation is dominating. In contrast, for *H ≪* 1, the response-*on* subpopulation contributes very little to the total mutant count.

Overall, in the best-case scenario and for the parameter regime considered here, *H ∼ 𝒪*(1) is sufficient for an accurate and precise estimate of the mutation-rate increase.

### 3.2 The impact of cell death

Our model accounts for changes in mutation and division rates upon induction of the stress response but neglects other potential consequences of the stress, such as cell death. Previous work showed that the occurrence of cell death, if neglected in the inference model, leads to overestimation of mutation rate [16]. Therefore, we asked whether neglecting cell death has a similar effect in our heterogeneous-response model. For this purpose, we simulate fluctuation assays under an extended model of heterogeneous stress responses with cell death. We consider the cases that (a) only response-*off* cells are affected by cell death, (b) only response-*on* cells are affected, or (c) all cells are affected equally, using parameter ranges of 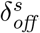, *δ*_*on*_ *∈* [0.0, 0.5] *h*^*−*1^.

Interestingly, we find that any biases in the estimated mutation-rate increase depend on which subpopulation is affected by cell death. If only response-*off* cells die, the mutationrate increase is overestimated for sufficiently large death rates (Figure 3A). On the other hand, if only response-*on* cells die, the mutation-rate increase is underestimated (Figure 3B). The estimation remains unbiased if both subpopulations are equally affected by cell death. However, the variation in the estimates increases for large death rates (Figure 3C). From the contribution of the response-*on* subpopulation to the mutant count given in equation 18, it can be seen that the effects of cell death in response-*off* and response-*on* cells partly cancel each other out. This result also holds for other parameter values (Supplementary Material Figure S2).

**Figure 3:**
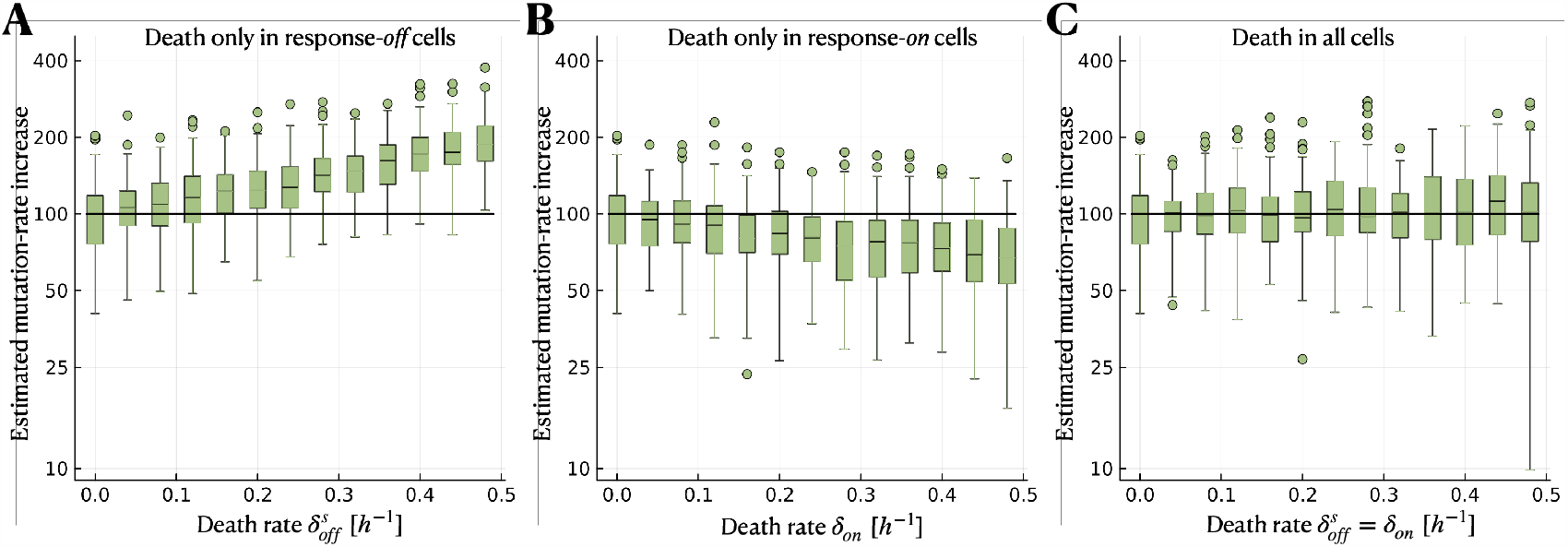
Cell death has limited impact on the estimation of the mutationrate increase. We simulate using the heterogeneous-response model (without differential mutant fitness and with zero division rate of response-*on* cells) but with cell death. However, we neglect cell death to infer *µ*_*off*_ and *µ*_*on*_. The black solid lines indicate the true mutation-rate increase used in the simulations. **A** Estimated mutation-rate increase when only response-*off* cells are affected by cell death, **B** when only response-*on* cells are affected by cell death, and **C** when all cells are affected by cell death equally. The parameter ranges used in the simulations are 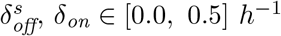, *δ*_*on*_ *∈* [0.0, 0.5] *h*^*−*1^ (with 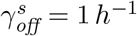).

We test another biologically realistic model deviation: the fitness of mutants differs from non-mutants during the growth phase (Supplementary Material Figure S3). We find that neglecting this effect in the inference leads to a slight underestimation of the mutation-rate increase when mutants have a fitness cost. On the other hand, if mutations bring a fitness advantage (for example, due to cross-resistance), the mutation-rate increase is slightly overestimated.

### 3.3 Estimation when response-*on* cells have a non-zero division rate

So far, we considered response-*on* cells not to divide at all, motivated by the SOS response. However, the division rate of response-*on* cells might be non-zero, especially if cells are exposed to a very low level of DNA damage (in the case of SOS) or for other stress responses. As a default setting, our method sets the relative fitness of response-*on* cells to zero (*r*_*on*_ = 0), but it also allows us to estimate *r*_*on*_ as an inference parameter. In the following, we evaluate the performance of our method when *r*_*on*_ *>* 0 and how this impacts the estimated mutation-rate increase.

We simulate under the heterogeneous-response model with a non-zero division rate of response-*on* cells, considering a parameter range of *γ*_*on*_ *∈* [0, 1] *h*^*−*1^ (with *γ*_*off*_ = 1 *h*^*−*1^). Note that the relative fitness of response-*on* cells *r*_*on*_ is equivalent to their division rate *γ*_*on*_ because we consider no cell death here. In the inference, we consider two different cases. In one case, we infer the mutation rates *µ*_*off*_ and *µ*_*on*_ and the relative fitness of response-*on* cells *r*_*on*_. Alternatively, we infer only *µ*_*off*_ and *µ*_*on*_ while setting *r*_*on*_ = 0. We estimate the mutation-rate increase 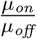 in both cases and compare it to the true value. In the first case, we also compare the estimated with the true *r*_*on*_.

We find that the estimation of the mutation-rate increase remains accurate for small to moderate relative fitness *r*_*on*_. For large *r*_*on*_ *→* 1, on the other hand, the mutation-rate increase is underestimated, yet more accurate and precise if *r*_*on*_ is also inferred (Figure 4A). However, the estimate of *r*_*on*_ itself is not precise and underestimated for large *r*_*on*_ (Figure 4B).

**Figure 4:**
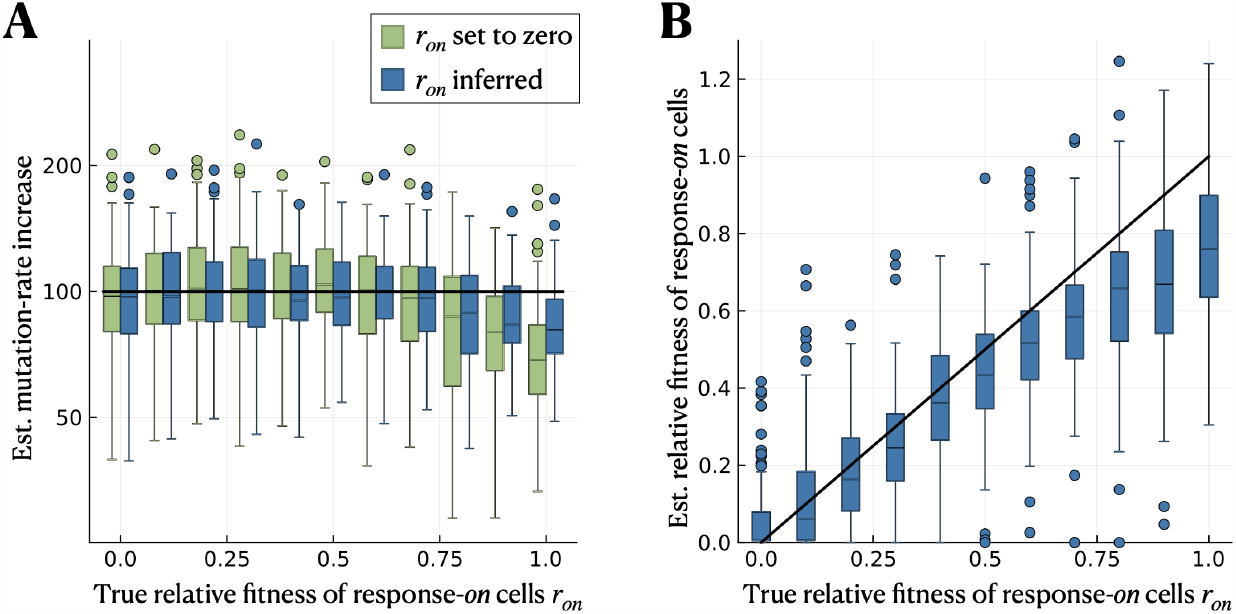
Estimation of the mutation-rate increase remains accurate when response-*on* cells have a small to moderate relative fitness. We simulate using the heterogeneous-response model (without cell death or differential mutant fitness) with *r*_*on*_ *≥* 0 being the relative fitness of response-*on* cells compared to response-*off* cells. We consider two cases for the inference: (i) setting *r*_*on*_ to zero and only inferring *µ*_*off*_ and *µ*_*on*_, and (ii) inferring *r*_*on*_ in addition to the two mutation rates. **A** Estimated mutation-rate increase 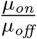 for both cases and a range of relative fitness of response-*on* cells. The solid black line indicates the true value of 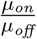. **B** Estimated compared to true relative fitness of response-*on* cells in inference case (ii). The parameter range used in the simulations is *γ*_*on*_ *∈* [0, 1] in *h*^*−*1^.

We also evaluate the performance when *r*_*on*_ is set to the true value in the inference. Interestingly, this increases the accuracy and precision of the estimate of 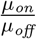 only slightly compared to when *r*_*on*_ is inferred (Supplementary Material Figure S4). The reason for this lies in the approximation (equation 5) made to derive the mutant count distribution, which is no longer valid for large *r*_*on*_.

### 3.4 Model selection between heterogeneous and homogeneous response

In many cases, it may not be known *a priori* whether the stress response is heterogeneously expressed across the population or whether, in contrast, all cells respond similarly. We want to determine whether distinguishing these two models is possible using mutant count data from fluctuation assays. To do so, we simulate fluctuation assays under the heterogeneous-response model for a range of relative fitness of response-*on* cells, *r*_*on*_. For the inference, we use both the heterogeneous- and homogeneous-response models and compare how well they fit the data. We use the same simulation data as in the previous section (parameter range *γ*_*on*_ *∈* [0, 1] *h*^*−*1^). However, we assume that the final population sizes 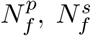 are known but not the fraction of the response-*on* subpopulation *f*_*on*_. Therefore, when using the heterogeneous-response model in the inference, we additionally infer *f*_*on*_.

We perform model selection between the heterogeneous-response and the homogeneous-response model using the AIC, which considers how well the models reproduce the data and penalises the number of model parameters. For the heterogeneous-response model, we consider two cases: (i) zero relative fitness of response-*on* cells (inference parameters: *µ*_*off*_, *µ*_*on*_ and *f*_*on*_) and (ii) non-zero relative fitness of response-*on* cells (inference parameters: *µ*_*off*_, *µ*_*on*_, *f*_*on*_ and *r*_*on*_). For the homogeneous-response model, we also consider two cases: (iii) without differential mutant fitness (inference parameters: *µ*^*p*^ and *µ*^*s*^) and (iv) with differential mutant fitness (inference parameters: *µ*^*p*^, *µ*^*s*^, *ρ*^*p*^ and *ρ*^*s*^). Then, we compare the estimated increase in population mean mutation rate from the best-fitting heterogeneous- and homogeneous-response models (given by 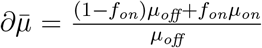 and 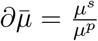, respectively) with the true value.

Comparing model fit, we find that the heterogeneous response model is selected only in a minority of cases (Figure 5A). The best case is when the relative fitness of response-*on* cells is small, where the heterogeneous-response model with *r*_*on*_ = 0 is selected for *≈*50% of the simulations. The homogeneous-response model with differential mutant fitness is selected in *≈*10% of simulations, while no model is clearly preferred for the remaining *≈*40%. For most simulations with intermediate values of *r*_*on*_, the heterogeneous-response model and the homogeneous-response model fit the simulated data comparably well, and no model can be selected. Eventually, for large *r*_*on*_, the homogeneous-response model without differential mutant fitness is selected in the majority of cases.

**Figure 5:**
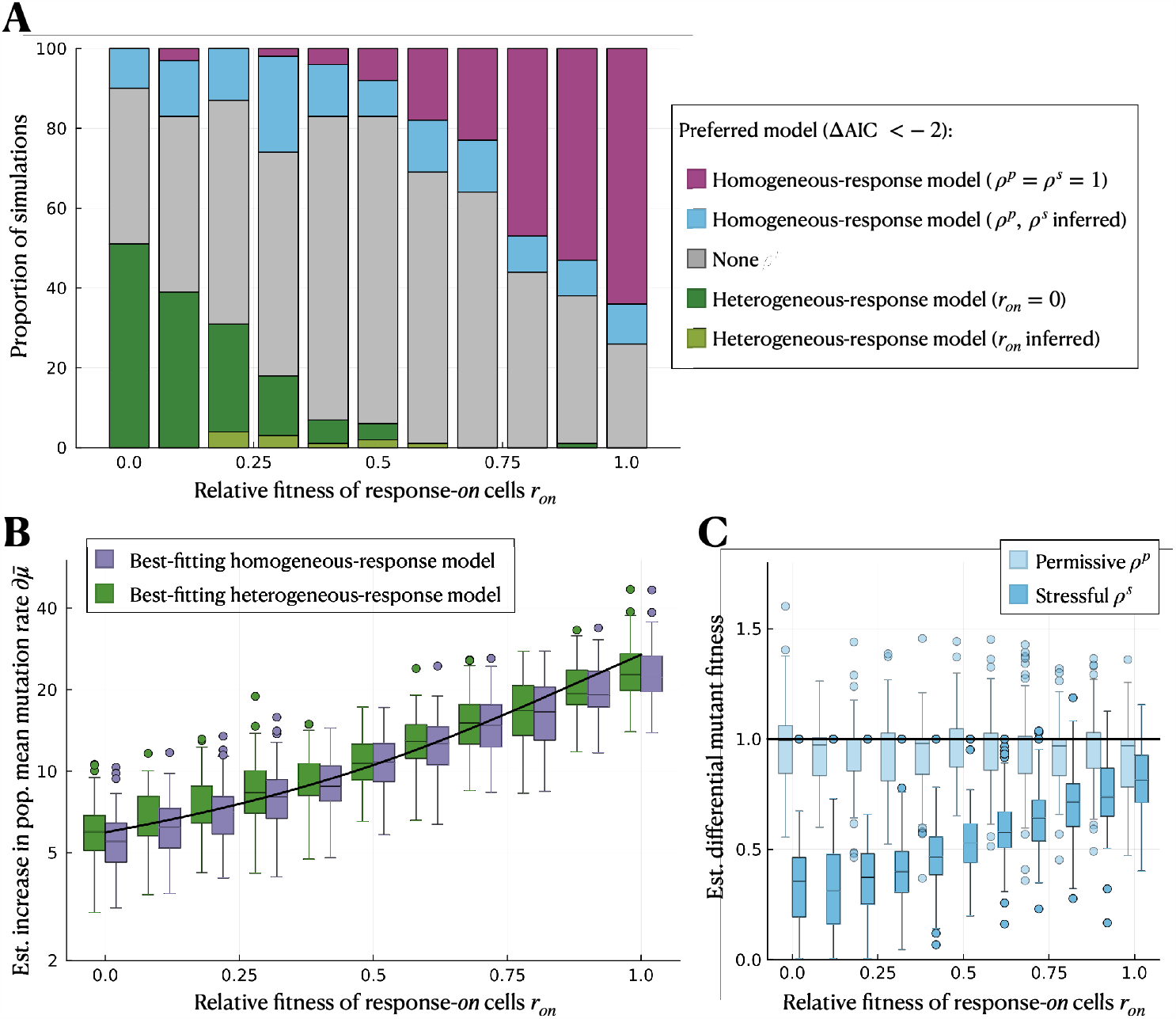
Heterogeneous-response and homogeneous-response models can reproduce mutant count data similarly well. We simulate under the heterogeneous-response model for a range of relative fitness of response-*on* cells, *r*_*on*_. In the inference, we use (i) the heterogeneous-response model with *r*_*on*_ = 0 (dark green) and (ii) the heterogeneousresponse model with *r*_*on*_ as an inference parameter (light green) to infer the mutation rates *µ*_*off*_ and *µ*_*on*_, the fraction of the response-*on* subpopulation *f*_*on*_ and, for (ii), additionally *r*_*on*_. We also use the homogeneous-response model (iii) without differential mutant fitness (*ρ*^*p*^ = *ρ*^*s*^ = 1; purple) and (iv) with differential mutant fitness (light blue) to infer the population-wide mutation rates under permissive and stressful conditions, *µ*^*p*^ and *µ*^*s*^, and, for (iv), additionally *ρ*^*p*^ and *ρ*^*s*^. **A** Model selection using the AIC. **B** Estimated increase in population mean mutation rate by the best-fitting heterogeneous-response model 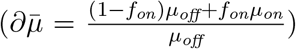 and the best-fitting homogeneous-response model 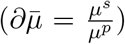. **C** Estimated differential mutant fitness in inference case (iv). The black lines indicate the true values. The parameter range used in the simulations is *γ*_*on*_ *∈* [0, 1] *h*^*−*1^.

Meanwhile, the estimated increase in population mean mutation rate, 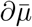, is quite accurate and precise over a wide range of *r*_*on*_, using either the heterogeneousor the homogeneousresponse model (Figure 5B). Generally, the homogeneous-response model estimates a slightly smaller value for 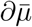 for small *r*_*on*_, the estimate is better when allowing for a differential mutant fitness (Supplementary Material Figure S6). In this case, the mutant fitness under the permissive condition is estimated to be around the true value, *ρ*^*p*^ *≈* 1. For the stressful condition, in contrast, an increasingly severe mutant cost is inferred as *r*_*on*_ decreases (Figure 5C). For intermediate *r*_*on*_, inference using the heterogeneous-response model improves the estimation of 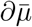 when inferring *r*_*on*_ instead of setting *r*_*on*_ = 0. (Supplementary Material Figure S6).

We also consider the homogeneous-response model with differential mutant fitness values constrained to be equal under stressful as under permissive conditions. The estimate of 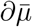 is less accurate using this model (Supplementary Material Figure S6). Moreover, for small *r*_*on*_, the heterogeneous-response model is preferred for most simulations (Supplementary Material Figure S5).

We repeat the analysis for a smaller mutation-rate increase,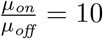, using unconstrained and then constrained mutant fitness for the homogeneous-response model. In this case, the heterogeneous-response model is selected only for a small fraction of simulations, but the estimation of the increase in population mean mutation rate remains accurate (Supplementary Material Figure S7 and S8).

## 4 Discussion

Since its introduction 80 years ago, the standard model behind the fluctuation assay has been extended numerous times to overcome limitations and make it more biologically realistic. Extensions particularly relevant for quantifying stress-induced mutagenesis include considering cell death [16, 37] and differential mutant fitness [38]. The latter can become important when resistance allowing growth on the selective plates also confers an advantage to the stressor (for example, due to cross-resistance) or when mutants carry a fitness cost. In this study, we addressed a so-far overlooked limitation of fluctuation analysis: heterogeneity in the expression of stress responses, which single-cell studies have recently demonstrated. Our population dynamic model considers response-*off* and response-*on* subpopulations, which allows us to estimate the relative increase in mutation rate associated with the induction of the stress response, 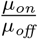.

We tested our estimation method with simulated mutant count data, which confirmed accurate and precise estimation of 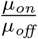 for sufficiently large mutation-rate heterogeneity 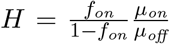 (Results section 3.1). *H* depends on the mutation-rate increase itself and the fraction of the response-*on* subpopulation *f*_*on*_. While 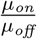 is inherent to the stress response, *f*_*on*_ could potentially be increased through experimental design. For example, increasing the antibiotic concentration has been shown to increase the rate of switching *on* the stress responses and thus the fraction of the response-*on* subpopulation [25]. Our results suggest that mutation rate estimates would be more accurate at higher antibiotic concentrations, all else being equal. Increasing antibiotic concentration could, however, also increase cell death. We neglect cell death in our inference, but we showed that our method is robust to this model deviation up to moderate death rates when cell death affects response-*off* and response-*on* subpopulations equally (Results section 3.2). Our estimation method also assumes the response-*on* cells are non- or slow-dividing, as in the SOS response. However, the estimated mutation-rate increase is robust to division rates up to *≈*75% of response-*off* cells and only marginally improved by inferring *r*_*on*_ rather than setting it to zero (Results section 3.3).

We used formal model selection with the AIC to evaluate whether a signal of mutation-rate heterogeneity can be detected from fluctuation assays alone. When response-*on* cells have low relative fitness compared to response-*off* cells (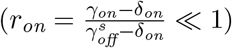), the heterogeneousresponse model is preferred over standard (homogeneous-response) models in about half of the simulated experiments (Results section 3.4). However, as *r*_*on*_ increases, homogeneousresponse models fit the data comparably well and eventually better at large *r*_*on*_ (due to the breakdown of the approximation made to derive the mutant count distribution). This implies that heterogeneity in stress responses may have been overlooked when using fluctuation assays, and these data should be complemented with additional experiments to support or rule out alternative explanations. For example, mutants arising in the fluctuation assay can be isolated to measure their fitness and check whether mutant fitness values estimated from the homogeneous-response model fit (*ρ*^*p*^ and *ρ*^*s*^ in permissive and stressful conditions, respectively) are reasonable. In particular, a large difference in estimated *ρ*^*p*^ and *ρ*^*s*^ may alternatively indicate the presence of a slowly-dividing and highly-mutating subpopulation (Figure 5C). Constraining *ρ*^*p*^ = *ρ*^*s*^, arguably a reasonable null model, increases the fraction of simulated experiments in which the heterogeneous model is preferred (Supplementary Material Figure S5).

If there is reason to suspect heterogeneity in the stress response, experimentalists can test this directly by engineering fluorescent reporters into the bacterial strain of interest and measuring the response on a single-cell level, e.g. by flow cytometry or microscopy. These experiments would provide an independent estimate of the fraction of response-*on* subpopulation to further constrain the heterogeneous response model. In reality, multiple factors causing deviation from the standard fluctuation assay model (e.g. heterogeneous responses, differential mutant fitness, and cell death) will likely operate simultaneously. Since it is not feasible to reliably estimate a large number of parameters from fluctuation assay data alone, separate experiments become important to decide which deviation(s) are most relevant to incorporate into the fluctuation analysis.

Interestingly, both the heterogeneousand the homogeneous-response models perform equally well in estimating the increase in population mean mutation rate (Results section 3.4). Therefore, neglecting heterogeneity in stress-induced mutagenesis has minimal effect on the estimation of increases in population mean mutation rate, and estimates from previous studies [7, 8, 9, 10, 11, 12, 13] are reliable, if interpreted as population-wide mutation rates. However, these studies may underestimate the true extent of mutagenesis associated with the expression of the stress response if it is only induced by a subpopulation of cells. Estimating not only the increase in *population mean* but also heterogeneity in mutation rate, as is possible with our method, could be important for parameterising evolutionary models, such as predictions of antibiotic resistance evolution. Theoretical modelling suggests that single-locus adaptation can be accurately captured by the population mean mutation rate, but cell-to-cell variation can speed up multi-locus adaptation [39]. However, this previous model did not incorporate any coupling of changes in mutation rate to changes in cell division or death rates, as would be expected in the case of stress responses. Therefore, an important direction for future work is to assess when the pleiotropic effects of realistic stress responses truly accelerate evolution.

Our approach to quantifying stress-induced mutagenesis assumes that the expression of the stress response is bimodal and can reasonably be modelled as either switched *off* or *on*. To a reasonable approximation, this expression pattern has been observed for the SOS response [25]. For some other stress responses, it might be more appropriate to model the expression as a continuous distribution. We expect, however, that this increase in model complexity would make parameter inference more challenging. We also neglect response-*on* cells switching the response *off* so long as the stress remains present during a fluctuation assay’s comparably short growth phase [25]. This assumption implies that the model cannot capture pulse-like mutagenesis as observed under DNA alkylation damage [21] and oxidative stress [40].

To be able to derive an analytical expression for the mutant count distribution, we make a series of approximations, the most important one being that cells with stress response switched *on* have a net growth rate (*γ*_*on*_ *−δ*_*on*_) much lower than that of *off* cells 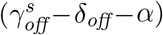 . For the SOS response, this approximation is valid, as induction of the response inhibits cell division. However, it might be violated for other stress responses, particularly if they protect cells from death, resulting in *δ*_*on*_ *< δ*_*off*_. In this case, our approximation is no longer valid, and therefore, parameter estimation using our method is expected to be less accurate.

In summary, we have presented and validated a new method for inferring stress-induced increases in mutation rate from fluctuation assays. Importantly, however, both a heterogeneous stress response and a homogeneous response with mutant fitness costs can generate similar patterns in fluctuation assay data and call for further experiments to distinguish these models. While both models can estimate the increase in population mean mutation rate, our new method of inferring heterogeneous mutation rates would be crucial for accurately characterising the mutagenic effects of stress responses and parameterising models of multi-locus adaptation. In future work, we aim to incorporate our new method into userfriendly tools for application to experimental data, similar to existing R packages [33, 41] and web tools [42, 32, 15, 43] for fluctuation analysis.

## Supporting information

Supplementary Material sections S1-S9

## Author contributions

LLJ: Conceptualization, Formal Analysis, Investigation, Methodology, Software, Validation, Visualization, Writing – Original Draft Preparation, Writing – Review & Editing \ MEK: Conceptualization, Funding Acquisition, Methodology, Supervision, Writing – Review & Editing HKA: Conceptualization, Funding Acquisition, Methodology, Project Administration, Supervision, Validation, Writing – Review & Editing

## Acknowledgements

The authors are grateful for helpful feedback on the mathematical results received from Tibor Antal and for the discussion and inspiration provided by the Alexander and El Karoui labs. This work was supported by the UKRI Biotechnology and Biological Sciences Research Council (BBSRC) grant number BB/T00875X/1 and a University of Edinburgh Principal’s Career Development PhD Scholarship to LLJ, a Wellcome Trust Investigator Award 205008/Z/16/Z to MEK, and a Royal Society University Research Fellowship URF/R1/191269 to HKA.

## Data Accessibility

The complete annotated documentation of the computational analyses of this study will be archived upon publication and is currently deposited at https://github.com/LucyL-J/Quantifying-SIM.

## Supplementary Material

**S1** Deterministic treatment of response-*on* non-mutants

**S2** Calculation of the duration of the growth phase

**S3** The impact of cell death

**S4** The impact of differential mutant fitness

**S5** Estimation when response-*on* cells have a known non-zero division rate

**S6** Model selection between heterogeneous response and homogeneous response with constrained mutant fitness

**S7** Estimation results from all heterogeneous- and homogeneous-response models

**S8** Model selection between heterogeneous and homogeneous response for smaller mutationrate increase

**S9** Non-identifiability of the heterogeneous-response model when the fraction of the response-*on* subpopulation is unknown

