## Supplementary Material sections S1-S9 for "Estimating Mutation Rates Under Heterogeneous Stress Responses"

November 25, 2023

[1]Institute of Ecology and Evolution, School of Biological Sciences, University of Edinburgh, UK

[2]Institute of Cell Biology, School of Biological Sciences, University of Edinburgh, UK

[3]Centre for Engineering Biology, University of Edinburgh, UK

*Corresponding authors

 (LLJ), (HKA)

### S1 Deterministic treatment of response-*on* non-mutants

In Model & Methods section 2.1, we derive the population sizes of the response-*off* and response-*on* non-mutants as

$$n_{off}(t) = n_{off}(0)\mathrm{e}^{(\gamma^s_{off}-\delta_{off}-\alpha)t} \tag{1}$$

$$n_{on}(t) = \frac{\alpha n_{off}(0)}{\gamma^s_{off}-\delta_{off}-\alpha-(\gamma_{on}-\delta_{on})}\left(\mathrm{e}^{(\gamma^s_{off}-\delta_{off}-\alpha)t}-\mathrm{e}^{(\gamma_{on}-\delta_{on})t}\right)+n_{on}(0)\mathrm{e}^{(\gamma_{on}-\delta_{on})t} \tag{2}$$

which assumes that the response-*on* subpopulation can be treated deterministically. However, for small initial population sizes $n_{on}(0)$, this assumption might not hold. Therefore, we test its validity using stochastic simulations; we simulate switching *on* of the response as a time-inhomogeneous Poisson process, the growth dynamics of the response-*on* subpopulation as a stochastic birth-death branching process until it reaches a size of $N = 10^4$ and according to equation 2 afterwards. Then, we compare the resulting population size with equation 2 at early and late time points: $t_1$, the expected time of the first mutation (with an exponential waiting time) in the response-*off* subpopulation, given by

$$t_1 = \frac{\Gamma\left(0, \frac{\mu_{off} n_{off}(0)}{\gamma^s_{off}-\delta_{off}-\alpha}\right)}{\gamma^s_{off}-\delta_{off}-\alpha}, \tag{3}$$

and $t_N$, the time when the response-*off* subpopulation reaches a size of $N = 10^9$, given by

$$t_N = \frac{\log\left(\frac{10^9}{n_{off}(0)}\right)}{\gamma^s_{off}-\delta_{off}-\alpha}. \tag{4}$$

For two initial population sizes of the response-*on* subpopulation, (i) $n_{on}(0) = 0$ and (ii) $n_{on}(0) = \frac{\alpha}{\gamma^s_{off}-\delta_{off}} \cdot n_{off}(0)$, which is a lower bound for the equilibrium fraction of the response-*on* subpopulation, we calculate the error relative to the median of $R = 100$ simulations and the coefficient of variation across the simulations (Figure S1). We find that both relative error and coefficient of variation are larger at the earlier than the later time point and that they increase with decreasing relative switching rate $\tilde{\alpha}$ and increasing relative fitness of response-*on* cells $r_{on}$. In general, equation 2 tends to slightly overestimate the size of the response-*on* subpopulation (indicated in red), but the error is with $< 1\,\%$ sufficiently small to justify our approximation.

#### S2 Calculation of the duration of the growth phase

In all simulations to test our inference method, we set the duration $t_f$ of the growth phase such that the expected number of *mutations*, $m$ (not *mutants*) equals one, by numerically solving the following equations:

$$\text{Permissive conditions: } \mathbb{E}[m] = \int_0^{t_f} \nu_{\mathit{off}} n_{\mathit{off}}(t)\,\mathrm{d}t \stackrel{!}{=} 1, \quad (\text{with } \alpha = \delta_{\mathit{off}} = 0) \tag{5}$$

$$\text{Stressful conditions: } \mathbb{E}[m] = \int_0^{t_f} \nu_{\mathit{off}} n_{\mathit{off}}(t) + \nu_{on} n_{on}(t)\,\mathrm{d}t \stackrel{!}{=} 1 \tag{6}$$

This way, the resulting number of resistant mutant colonies on each selective plate is similar across the considered parameter ranges and usually within an experimentally countable range of zero to a couple hundred. For example, the duration of simulated fluctuation assays under permissive conditions is set to $t_f \approx 9.21\,h$.

#### S3 The impact of cell death

In Results section 3.2, we show that cell death impacts the estimation of the mutation-rate increase depending on which subpopulation is affected by death. However, estimates remain largely unbiased when all cells are affected equally, although their variation increases. This

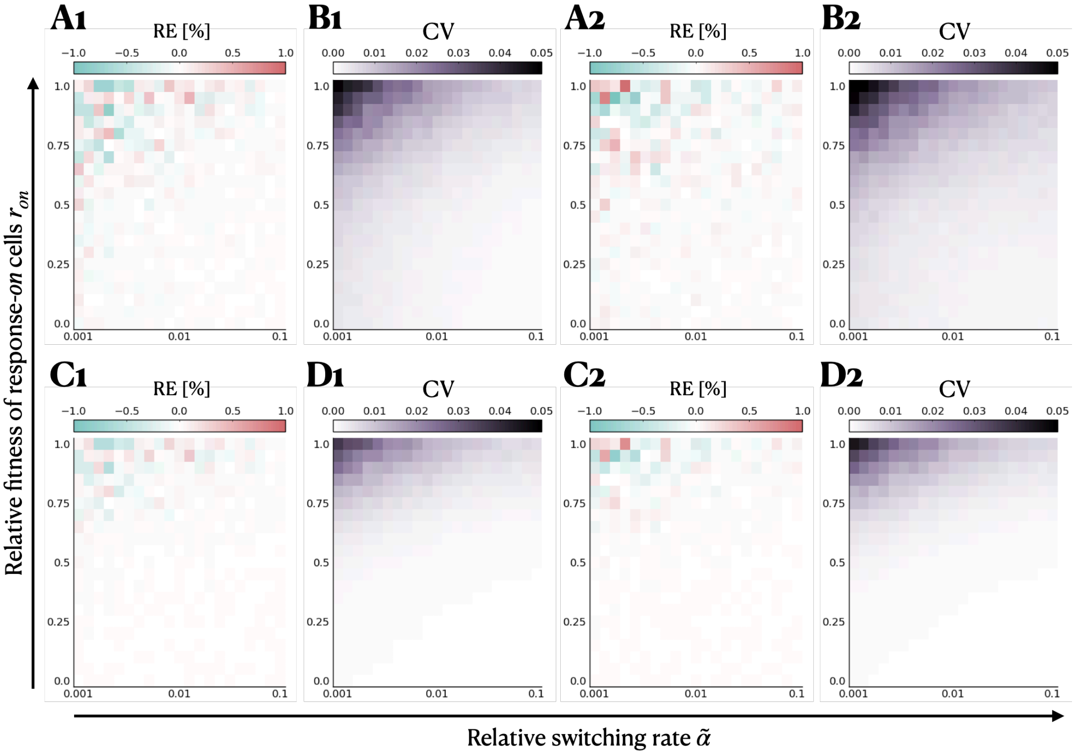


Figure S1: **Response-*on* non-mutants can be treated deterministically.** Deterministic compared to stochastic dynamics of response-*on* non-mutants for initial population sizes of the response-*on* subpopulation of (**1**) $n_{on}(0) = 0$ and (**2**) $n_{on}(0) = \frac{\alpha}{\gamma^s_{off} - \delta_{off}} \cdot n_{off}(0)$. We simulate switching *on* of the response as a time-inhomogeneous Poisson process and the dynamics of response-*on* non-mutants as stochastic birth-death processes ($R = 100$ simulation runs). Then, we compare the resulting population sizes with equation 2 at early and late time points. **A** Relative error of the median in % at $t_1$, **B** coefficient of variation across the simulations at $t_1$, **C** relative error of the median in % at $t_N$ and **D** coefficient of variation across the simulations at $t_N$. The parameters used in the simulations are, $\alpha \in [0.001,\ 0.1]\ h^{-1}$, $\gamma_{on} \in [0.0,\ 1.0]\ h^{-1}$.

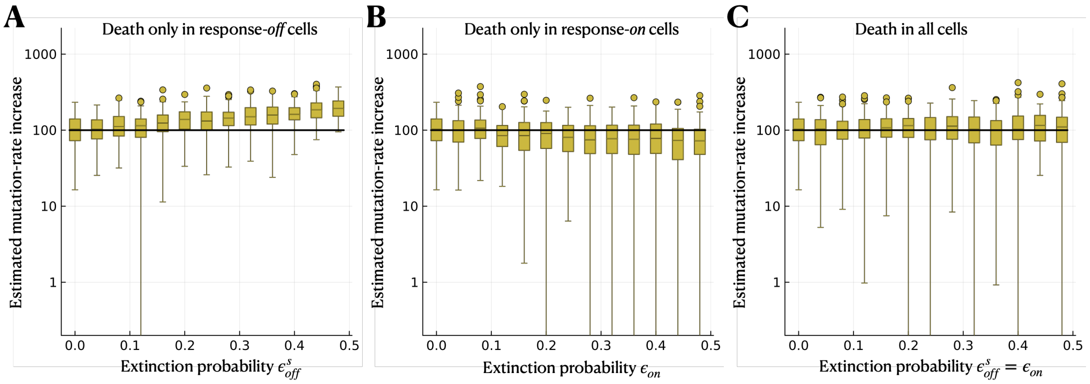


Figure S2: **Cell death has limited impact on the estimation of the mutation-rate increase.** Here, we repeat the analysis in Results Figure 3 but with a lower switching rate, $\alpha$. We simulate using the heterogeneous-response model extended by cell death but neglect cell death in the model used to infer $\mu_{off}$ and $\mu_{on}$. The black solid lines indicate the true mutation-rate increase used in the simulations. **A** Estimated mutation-rate increase when only response-*off* cells are affected by cell death, **B** when only response-*on* cells are affected by cell death and **C** when all cells are affected by cell death equally. The parameter range used in the simulations is $\alpha = 0.01\ h^{-1}$, $\delta_{off} \in [0.0,\ 0.5]\ h^{-1}$, $\delta_{on} \in [0.0,\ 0.5]\ h^{-1}$.

result is not an artefact of the specific parameters used in Results section 3.2: it also holds for different values of the relative switching rate, here shown with $\tilde{\alpha} = 0.01$ (Figure S2) instead of $\tilde{\alpha} = 0.05$ (Results Figure 3). A possible explanation for this result is that death in response-*off* cells causes an overestimation of the per-division mutation rate $\mu_{on}$ because, as pointed out in [1], the number of cell divisions to reach the final population sizes is underestimated. At the same time, death in response-*on* cells can lead to the extinction of response-*on* mutant lineages, causing an underestimation of $\mu_{on}$. These two effects counteract each other, leading to an unbiased estimate overall. The estimation of $\mu_{off}$ is not noticeably impacted by cell death due to the joint inference under stressful and permissive conditions, as for the latter, there is no cell death.

### S4 The impact of differential mutant fitness

In addition to cell death, we test the robustness of our method to differential mutant fitness, which is neglected in the inference. For this purpose, we simulate fluctuation assays under an extended model of heterogeneous stress responses with differential mutant fitness using a parameter range of $\rho_{off} := \rho^p_{off} = \rho^s_{off} \in [0.0,\ 1.5]$. Note that we consider a differential mutant fitness only in response-*off* cells as the response-*on* cells have a zero division rate in this set of simulations and are, therefore, unaffected by a differential mutant fitness.

From the resulting mutant count data, we estimate the mutation-rate increase $\frac{\mu_{on}}{\mu_{off}}$ and compare it with the true value to determine any biases caused by neglecting the differential mutant fitness in the inference (Figure S3). We find that neglecting this effect in the inference leads to a slight overestimation of the mutation-rate increase when mutants have a fitness cost ($\rho_{off} < 1$). On the other hand, if mutations bring a fitness advantage ($\rho_{off} > 1$), the mutation-rate increase is slightly underestimated. Fitness advantages could arise, for example, because the same antibiotic is used as a stressor and on the selective plates or because two different antibiotics are used, but mutants are cross-resistant [2]. However, the bias remains small (relative error of the median $< 12\,\%$) across the tested parameter range.

### S5 Estimation when response-*on* cells have a known non-zero division rate

In Results section 3.3, we evaluate the performance of our method in estimating the mutation-rate increase when response-*on* cells have a non-zero division rate for the cases that (i) the non-zero division rate is neglected in the inference (setting $r_{on} = 0$) and (ii)

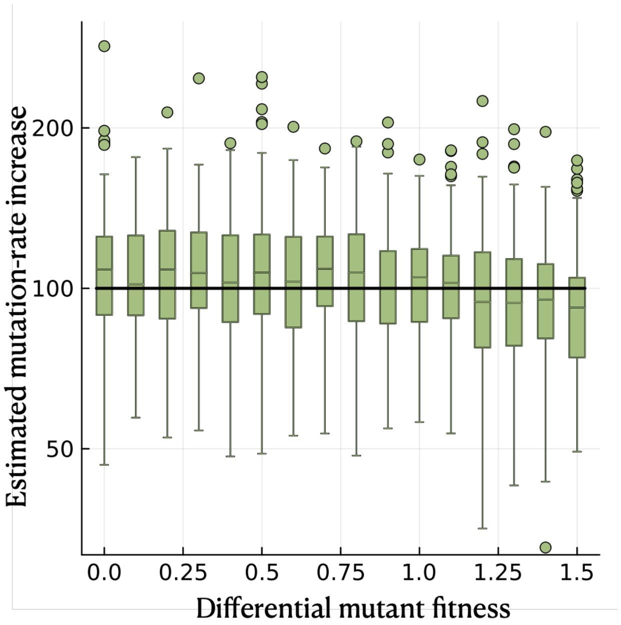


Figure S3: **Differential mutant fitness has minimal impact on the estimation of the mutation-rate increase.** We simulate using the heterogeneous-response model with mutants having a different division rate than non-mutants but neglect this effect in the model used to infer $\mu_{off}$ and $\mu_{on}$. The black solid line indicates the true mutation-rate increase. The parameter range used in the simulations is $\rho^p_{off} = \rho^s_{off} \in [0.0,\ 1.5]$.

$r_{on} > 0$ is inferred additionally. Here, we consider a third case: $r_{on}$ is set to the true value, which could, for example, be measured in microfluidics experiments using time-lapse microscopy and image analysis to estimate the division rate of cells identified as response-*on*. Interestingly, we find that setting $r_{on}$ to the true value hardly improves the estimation of the mutation-rate increase (Figure S4). The reason for this lies in the approximation of the size of the response-*on* subpopulation

$$\hat{n}_{on}(t) \approx \frac{\alpha n_{\mathit{off}}(0)}{\gamma^s_{\mathit{off}} - \delta_{\mathit{off}} - \alpha - (\gamma_{on} - \delta_{on})} \mathrm{e}^{(\gamma^s_{\mathit{off}} - \delta_{\mathit{off}} - \alpha)t}. \quad (7)$$

which assumes that $\gamma_{on} - \delta_{on} \ll \gamma^s_{\mathit{off}} - \delta_{\mathit{off}} - \alpha$ and is no longer valid for large $r_{on} = \frac{\gamma_{on} - \delta_{on}}{\gamma^s_{\mathit{off}} - \delta_{\mathit{off}}} \to 1$. Therefore, since we derive the mutant count distribution based on equation 7, setting $r_{on}$ to the true value still results in a biased estimate. This suggests that, given the available inference method, obtaining a precise estimate of $r_{on}$ is not worthwhile besides validating that it is small.

### S6 Model selection between heterogeneous response and homogeneous response with constrained mutant fitness

In Results section 3.4, we simulate under the heterogeneous-response model for a range of relative fitness of response-*on* cells ($r_{on}$) and perform model selection between (i) the heterogeneous-response model with zero fitness of response-*on* cells (setting $r_{on} = 0$), (ii) the heterogeneous-response model with $r_{on}$ inferred in addition to the mutation rates and the fraction of response-*on* cells, (iii) the homogeneous-response model without differential mutant fitness (setting $\rho^p = \rho^s = 1$), and (iv) the homogeneous-response model with differential mutant fitness, i.e. $\rho^p$ and $\rho^s$ inferred in addition to the mutation rates. Moreover, we evaluate the models' performance in estimating the increase in population mean muta-

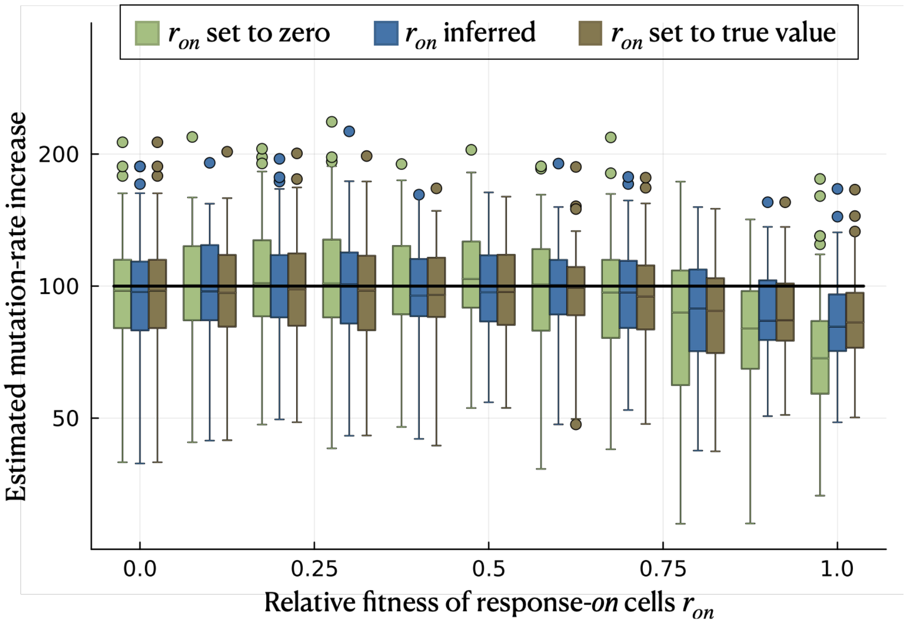


Figure S4: **Setting the relative fitness of response-*on* cells to the true value only marginally improves the estimation of the mutation-rate increase.** We simulate using the heterogeneous-response model with $r_{on} \geq 0$ being the relative fitness of response-*on* cells compared to response-*off* cells. We consider three cases for the inference: (i) setting $r_{on}$ to zero and only inferring $\mu_{off}$ and $\mu_{on}$, (ii) inferring $r_{on}$ in addition to the two mutation rates, and (iii) setting $r_{on}$ to its true value and only inferring $\mu_{off}$ and $\mu_{on}$. The solid black line indicates the true value of $\frac{\mu_{on}}{\mu_{off}}$. The parameter range used in the simulations is $\gamma_{on} \in [0.0,\ 1.0]\ h^{-1}$.

tion rate. Here, we repeat this analysis, but for model (iv), we constrain the differential mutant fitness to be equal under stressful and permissive conditions, $\rho^p = \rho^s$.

Comparing model fit, we find that the heterogeneous-response model with $r_{on} = 0$ is selected in most cases when the relative fitness of response-*on* cells $r_{on}$ is small (Figure S5A). The heterogeneous-response model with $r_{on}$ inferred is selected more often (up to $\sim 30\%$) for intermediate values of $r_{on}$. For large values of $r_{on}$, on the other hand, the homogeneous-response model without differential mutant fitness is selected in most cases. Similarly, the number of simulations for which no model is preferred is highest ($\sim 40\%$) for intermediate $r_{on}$. The homogeneous-response model with constrained differential mutant fitness is selected in only a few simulations, with the highest percentage of $\sim 20\%$ for large $r_{on}$. Over the whole parameter range, the heterogeneous-response model is selected more often compared to when the mutant fitness is not constrained to be equal under stressful and permissive conditions (Results section 3.4).

Moreover, we find that constraining the differential mutant fitness results in an underestimation of the increase in population mean mutation rate $\Delta\bar{\mu}$, especially for small $r_{on}$ (Figure S5), as the inference of the constrained differential mutant fitness does not improve the estimate compared to setting $\rho^p = \rho^s = 1$ (Figure S6A2). The differential mutant fitness itself ($\rho^p = \rho^s$) is estimated to be increasingly severe when the true relative fitness of response-*on* cells used in the simulations is smaller (Figure S5C).

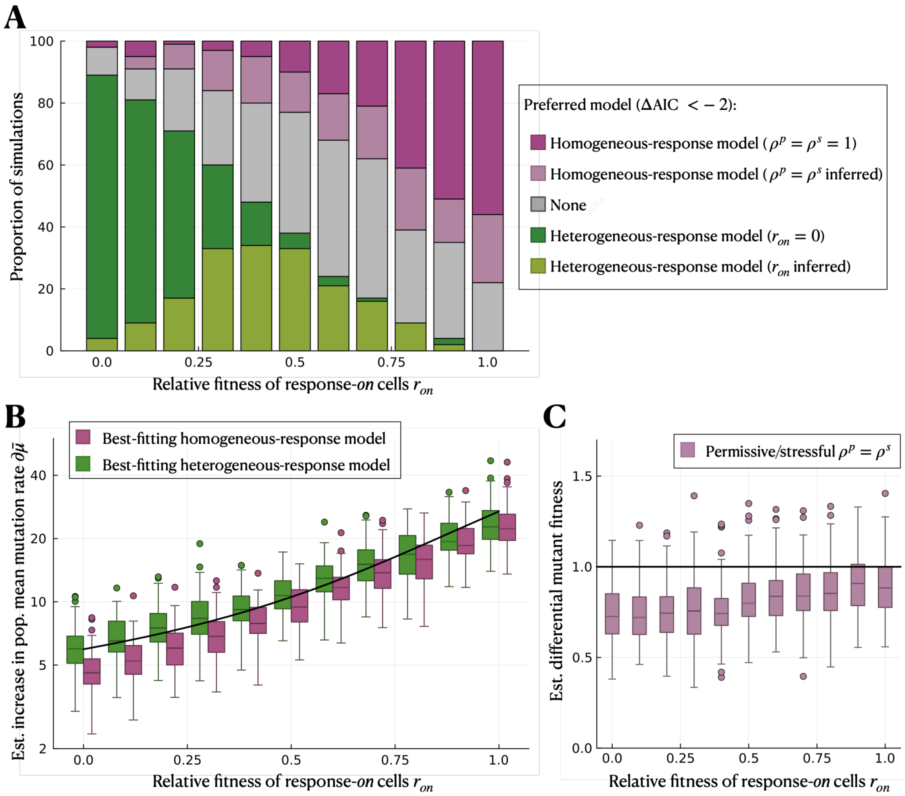


Figure S5: **Constraining mutant fitness means that the heterogeneous-response model is selected more often.** We simulate using the heterogeneous-response model for a range of relative fitness of response-*on* cells, $r_{on}$. In the inference, we use the heterogeneous-response model (i) with $r_{on} = 0$ (dark green) and (ii) with $r_{on}$ as an inference parameter (light green), as well as the homogeneous-response model (iii) without differential mutant fitness ($\rho^p = \rho^s = 1$; purple) and (iv) with differential mutant fitness constrained to be equal under permissive and stressful conditions ($\rho^p = \rho^s$; violet). **A** Model selection using the AIC. **B** Estimated increase in population mean mutation rate by the best-fitting heterogeneous- and homogeneous-response models (given by $\frac{(1-f_{on})\mu_{off}+f_{on}\mu_{on}}{\mu_{off}}$ and $\frac{\mu^s}{\mu^p}$, respectively). The black line gives the true value. **C** Estimated constrained differential mutant fitness in inference case (iv). The solid black line indicates the true value. The parameter range used in the simulations is $\gamma_{on} \in [0,\ 1]\ h^{-1}$.

### S7 Estimation results from all heterogeneous- and homogeneous-response models

In Results section 3.4, we show the estimated increase in the population mean mutation rate of the best-fitting heterogeneous- and homogeneous-response models. Here, we individually show the estimated parameters of all model versions (Figure S6).

### S8 Model selection between heterogeneous and homogeneous response for smaller mutation-rate increase

In Results section 3.4 and Supplementary Material section S6 above, we perform model selection between heterogeneous and homogeneous response using simulated data where we set the true mutation-rate increase to $\frac{\mu_{on}}{\mu_{off}} = 100$. Here, we repeat this analysis for a mutation-rate ratio of $\frac{\mu_{on}}{\mu_{off}} = 10$. We find that the number of simulations in which the heterogeneous-response model is selected drops, reaching at most $\sim 20\%$ for small $r_{on}$ (Figure S7A and S8A). In the remaining cases, either no model is preferred (more commonly for small $r_{on}$), or the homogeneous-response model without differential fitness is selected (more commonly for large $r_{on}$). The homogeneous-response models with inferred mutant fitness are selected in only a few cases, and they infer a small mutant cost over the whole range of $r_{on}$ (Figure S7C and SS8C). Both heterogeneous- and homogeneous-response models remain able to infer the increase in population mean mutation rate reasonably accurately, though again with slight underestimation in the homogeneous case (Figure S7B and SS8B).

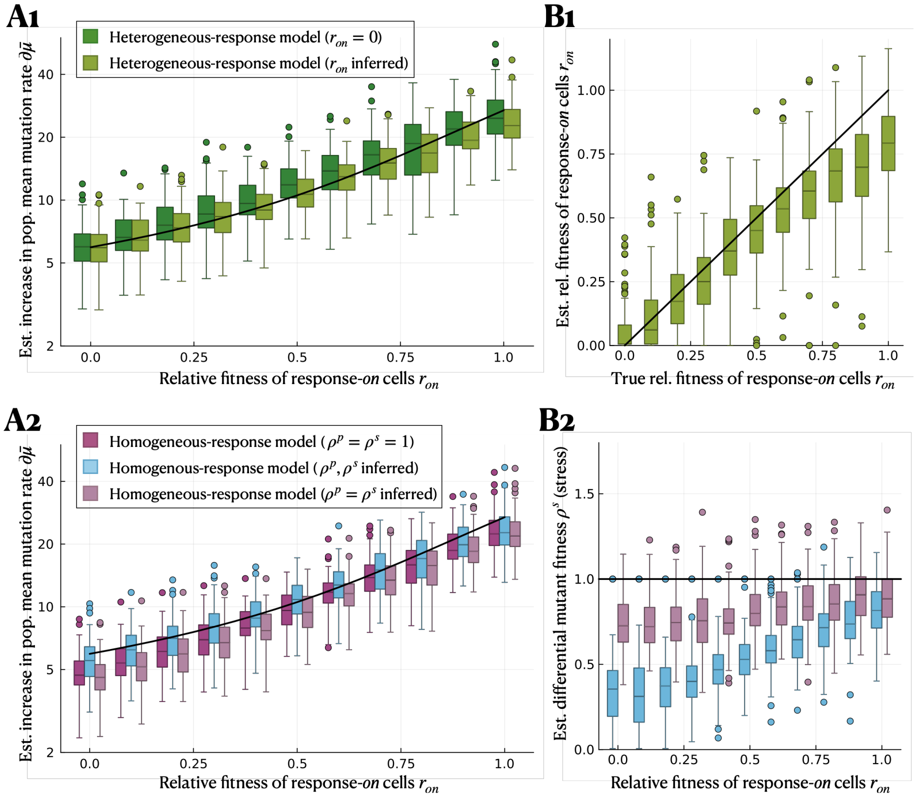


Figure S6: **Inferring additional model parameters improves the estimation of the mutation-rate increase.** Estimated parameters (**A**, increase in population mean mutation rate; **B** additional parameters) of **1** the heterogeneous-response model with $r_{on} = 0$ (dark green) versus $r_{on}$ inferred (light green), **2** the homogeneous-response model with $\rho^p = \rho^s = 1$ (purple) versus $\rho^p$ and $\rho^s$ inferred individually (light blue) or jointly ($\rho^p = \rho^s$) (violet). The same simulated data is used in Figure S5.

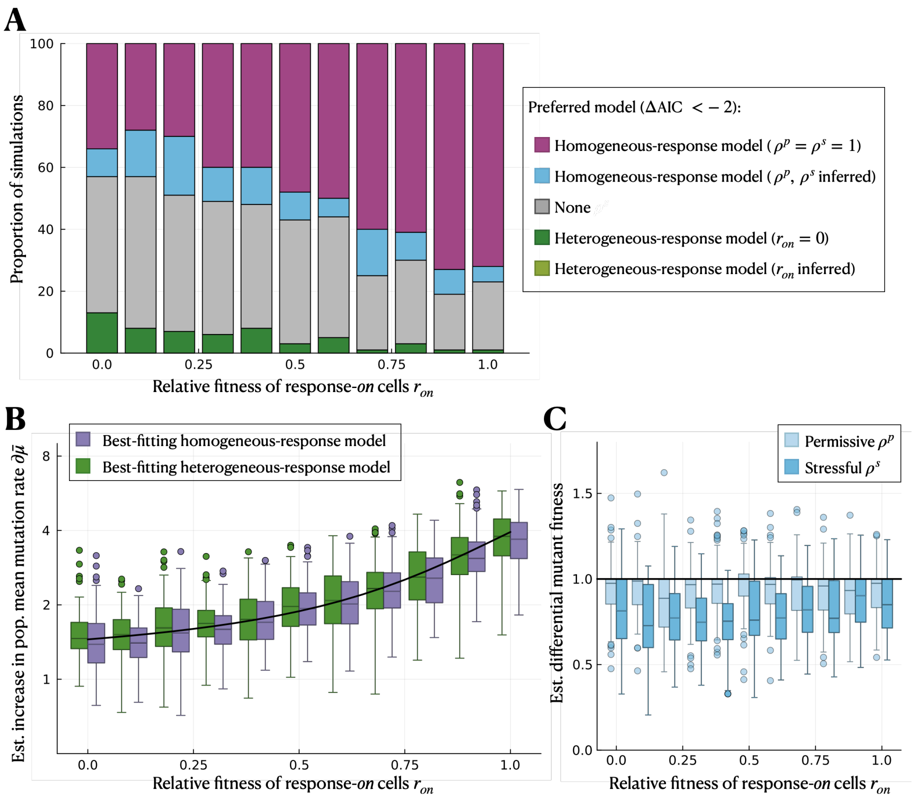


Figure S7: **The heterogeneous-response model is selected less often when the mutation-rate increase is smaller, compared to a homogeneous model with unconstrained mutant fitness.** We simulate using the heterogeneous-response model for a range of relative fitness of response-*on* cells, $r_{on}$, and with a true mutation-rate ratio of $\frac{\mu_{on}}{\mu_{off}} = 10$; and infer using the same models as in Results section 3.4. **A** Model selection using the AIC. **B** Estimated increase in population mean mutation rate by the best-fitting heterogeneous/homogeneous-response model (given by $\frac{(1-f_{on})\mu_{off}+f_{on}\mu_{on}}{\mu_{off}}$ and $\frac{\mu^s}{\mu^p}$, respectively). **C** Estimated differential mutant fitness under permissive and stressful conditions in inference case (iv). The solid black lines indicate the true value. The parameters used in the simulations are $\nu_{on} = 10^{-7}\ h^{-1}$ and $\gamma_{on} \in [0,\ 1]\ h^{-1}$.

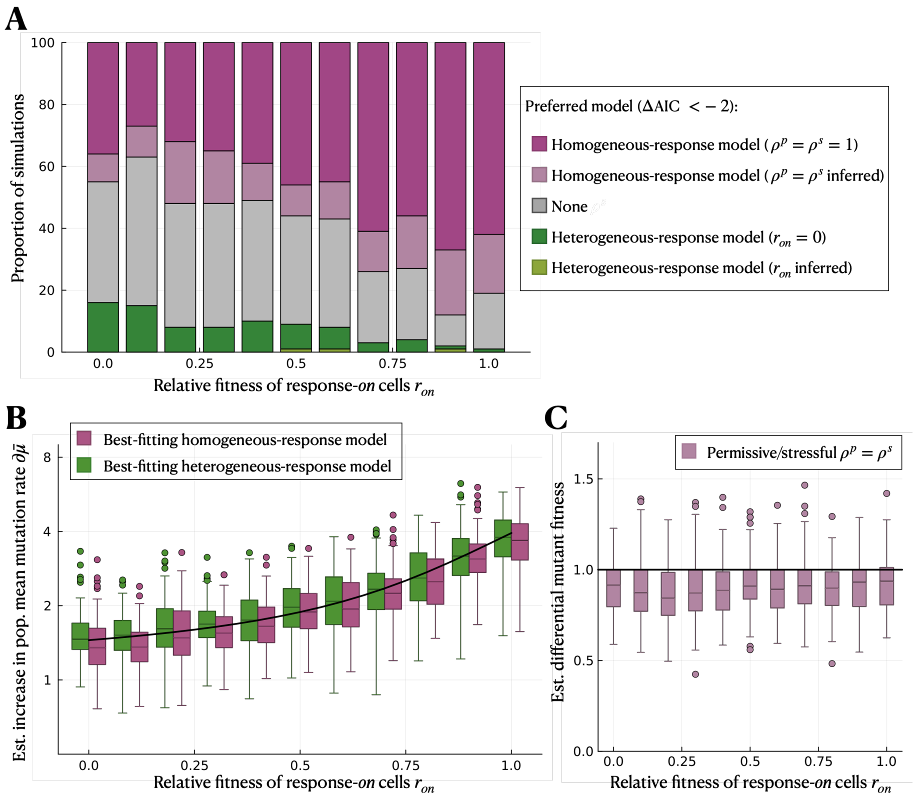


Figure S8: **The heterogeneous-response model is selected less often when the mutation-rate increase is smaller, compared to a homogeneous model with constrained mutant fitness.** We consider the same simulated data as in Figure S7 and, for the inference, the same models as in Figure S5. **A** Model selection, **B** estimated increase in population mean mutation rate and **C** estimated differential mutant fitness.

### S9 Non-identifiability of the heterogeneous-response model when the fraction of the response-*on* subpopulation is unknown

In Results section 3.4, we assume that the fraction of response-*on* cells ($f_{on}$) is unknown in the inference using the heterogeneous-response model. In this case, the heterogeneous-response model is non-identifiable because, for small relative fitness of response-*on* cells, the probability generating function of the mutant count distribution (Model & Methods equations 17, 18) reduces to

$$G^s_{\text{het}}(z) \overset{r_{on}\to 0}{=} \exp\left[\mu_{\mathit{off}} N^s_f (1-z)\left(\frac{\log(1-z)}{z} - \frac{\mu_{on}}{\mu_{\mathit{off}}}\frac{f_{on}}{1-f_{on}}\right)\right], \tag{8}$$

and for large $r_{on}$ to

$$G^s_{\text{het}}(z) \overset{r_{on}\to 1}{=} \exp\left[(\mu_{\mathit{off}}(1-f_{on}) + \mu_{on} f_{on}) N^s_f \frac{(1-z)\log(1-z)}{z}\right]. \tag{9}$$

In both cases, $\mu_{on}$ and $f_{on}$ appear only as a product, implying that they effectively cannot be inferred individually. We find that estimating these parameters separately is heavily biased, with $\mu_{on}$ and hence the mutation-rate increase being overestimated and $f_{on}$ underestimated. On the other hand, estimation of the mutation-rate heterogeneity, $H = \frac{\mu_{on}}{\mu_{\mathit{off}}}\frac{f_{on}}{1-f_{on}}$, is precise and accurate allowing for unbiased estimation of the increase in population mean mutation rate $\frac{(1-f_{on})\mu_{\mathit{off}} + f_{on}\mu_{on}}{\mu_{\mathit{off}}}$ (Figure S6). Note that the estimation of $\mu_{\mathit{off}}$ remains accurate due to the additional information from the fluctuation assay under permissive conditions.

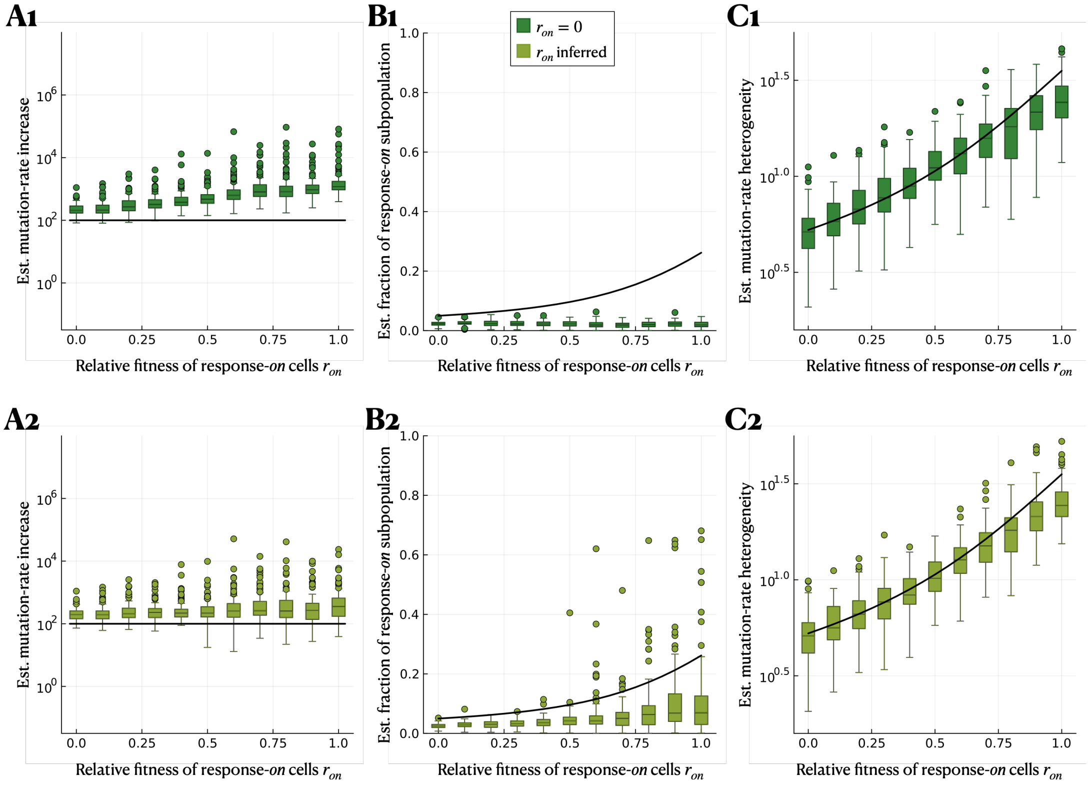


Figure S9: **The heterogeneous-response model is non-identifiable when the fraction of the response-*on* subpopulation ($f_{on}$) is unknown.** We simulate using the heterogeneous-response model for a range of relative fitness of response-*on* cells, $r_{on}$. We infer under the heterogeneous-response model with **1** setting $r_{on} = 0$ and **2** inferring $r_{on}$. **A** Estimated mutation-rate increase, $\frac{\mu_{on}}{\mu_{off}}$, and **B** estimated fraction of response-*on* subpopulation, $f_{on}$. The true values are indicated in solid black lines. The parameter range used in the simulations is $\gamma_{on} \in [0.0,\ 1.0]\ h^{-1}$.
